# Dimensionality reduction by UMAP reinforces sample heterogeneity analysis in bulk transcriptomic data

**DOI:** 10.1101/2021.01.12.426467

**Authors:** Yang Yang, Hongjian Sun, Yu Zhang, Tiefu Zhang, Jialei Gong, Yunbo Wei, Yong-Gang Duan, Minglei Shu, Yuchen Yang, Di Wu, Di Yu

## Abstract

Transcriptome profiling and differential gene expression constitute a ubiquitous tool in biomedical research and clinical application. Linear dimensionality reduction methods especially principal component analysis (PCA) are widely used in detecting sample-to-sample heterogeneity in bulk transcriptomic datasets so that appropriate analytic methods can be used to correct batch effects, remove outliers and distinguish subgroups. In response to the challenge in analysing transcriptomic datasets with large sample size such as single-cell RNA-sequencing (scRNA-seq), non-linear dimensionality reduction methods were developed. t-distributed stochastic neighbour embedding (t-SNE) and uniform manifold approximation and projection (UMAP) show the advantage of preserving local information among samples and enable effective identification of heterogeneity and efficient organisation of clusters in scRNA-seq analysis. However, the utility of t-SNE and UMAP in bulk transcriptomic analysis has not been carefully examined. Therefore, we compared major dimensionality reduction methods (linear: PCA; nonlinear: multidimensional scaling (MDS), t-SNE, and UMAP) in analysing 71 bulk transcriptomic datasets with large sample sizes. UMAP was found superior in preserving sample level neighbourhood information and maintaining clustering accuracy, thus conspicuously differentiating batch effects, identifying pre-defined biological groups and revealing in-depth clustering structures. We further verified that new clustering structures visualised by UMAP were associated with biological features and clinical meaning. Therefore, we recommend the adoption of UMAP in visualising and analysing of sizable bulk transcriptomic datasets.

## Introduction

Bulk transcriptome profiling quantifies the transcripts in a given biological sample, achieved by technologies including microarray [1, 2] and RNA sequencing (RNA-seq) [3, 4]. This tool is ubiquitously adopted in modern biomedical research and application to reveal unique features of gene expression for specific cell or tissue type and biological process. The principal task of bulk transcriptome profiling is to analyse differential gene expression (DGE) of samples between biological groups. When statistically modelling DGE, an implicit assumption is that data of individual samples within a given group are relatively homogeneous. For instance, to investigate the biomarker for a certain disease, the group comparison between patient and healthy control cohorts presumes that the biological characteristics of individual patients are largely indistinguishable when compared to healthy controls, and vice versa. However, there exists heterogeneity within a group, which can lie in samples’ distinct biological states. For example, patients with systemic lupus erythematosus (SLE) show distinct disease activities and can be classified based on the levels of disease activity index [5]. Other heterogeneity can result from different sample preparation or processing conditions, often referred to as batch effects [6, 7]. Therefore, it is crucial to scrutinise sample-to-sample heterogeneity within groups so that subgroups or outliers can be identified. Only with such information, appropriate analytic methods can be used to correct batch effects, remove outliers and distinguish subgroups. In contrast, DGE analysis simply in given groups without the knowledge of sample-to-sample heterogeneity within groups can often lead to biased or even wrong conclusion.

To detect among-sample heterogeneity in bulk transcriptome profiling, individual samples are visualised in embedded space by dimensionality reduction methods. Principal component analysis (PCA, [8]) and multidimensional scaling (MDS, [9]) have been thoroughly exploited to obtain an overview of sample relationship in a low-dimensional space [10-13]. Both methods succeeded in visualising biological or technical variation among samples by uncovering the overall structure of the sample-to-sample relationship, which represents the key information of among-sample heterogeneity.

Since 2009 [14], the new era of characterising transcriptome at single-cell level has arrived. Numerous single-cell RNA sequencing (scRNA-seq) technologies enable simultaneous profiling of thousands of cells’ transcriptomes in a given sample so that the analysis of population heterogeneity can identify complex compositions, reveal rare cell populations, detect differentially expressed genes between multiple cell populations or between samples for cell types, uncover cell differentiation trajectories, and so force [15, 16]. However, PCA and MDS show inefficient performance for dimensionality reduction of scRNA-seq data while two non-linear methods, *t*-distributed stochastic neighbour embedding (*t*-SNE) [17] and uniform manifold approximation and projection (UMAP) [18, 19] exhibit better capability due to the advantage of maintaining cell-to-cell neighbour information and visualising local structure. Compared to t-SNE, UMAP can not only distinguish neighbouring clusters but also retain the global structure in scRNA-seq data analysis [18, 19].

The continuous improvement and invention of sequencing platforms has hugely improved the efficiency and throughput of DNA sequencing and resulted in a dramatic reduction in costs, which enable to generate a large number of samples and datasets of bulk transcriptome profiling. For example, the landmark cancer genomics program – The Cancer Genome Atlas (TCGA) has profiled over 20,000 primary cancer and matched normal samples spanning 33 cancer types and generated over 2.5 petabytes of genomic, epigenomic, transcriptomic, and proteomic data [20-22]. While PCA remains as the mainstream tool recommended detecting among-sample heterogeneity in bulk transcriptome profiling, such as by TCGA Batch Effects [6, 7], we hypothesis that, for datasets with large sample sizes, local structure of sample-to-sample relationship becomes more prominent for sample heterogeneity analysis. Therefore, non-linear methods t-SNE and UMAP might outperform PCA and MDS.

In this study, we visually and quantitatively compared the capabilities of PCA, MDS, t-SNE, and UMAP in heterogeneity exploration of bulk transcriptome profiling. By visualising and interpreting 71 sizeable datasets of bulk transcriptome profiling, we found that UMAP was superior in preserving sample level neighbourhood information and maintaining clustering accuracy, thus conspicuously differentiating batch effects, identifying pre-defined biological groups and identifying new clustering structures associated with biological features and clinical meaning.

## Result

### Overview of the evaluation

The bulk-transcriptome profiling datasets were collected from the Gene Expression Omnibus (GEO) database within past five years (**Table S1**). To minimize the cell type effects interacting with our results which are usually strong and very easy to be identified, we only chose the datasets of human samples from peripheral blood mononuclear cells (PBMCs) or whole blood for bulk transcriptome analysis, that are among most frequent cell populations. Datasets with the size less than 100 samples were excluded in order to generate observable and meaningful clusters. The collection covered a diverse range of biomedical research including the investigations on disease features such as SLE [23-29] and influenza infection [30-32], and the evaluation on interventions such as therapies and vaccination [33-37].

The research design flowchart is shown in **Figure 1a**. Among a total collection of 71 datasets based on the above procedure, there were 41 datasets revealing clustering structures in plots of two-dimensional embedding space by the dimensionality reduction methods PCA, MDS, t-SNE and UMAP. UMAP reported all clustering (41/71) and, together with t-SNE (37/71), performed significantly better than PCA (11/71) and MDS (13/71) (**Figure 1b**). The 41 datasets were classified into three categories by incorporating available features (**Figure 1b**). As in **Figure 1b**, three plots in the two-dimensional embedding space from the dimension reduction methods showed clusters related to batches (batch effect) described in studies for these datasets while 9 plots showed clusters related to biological groups designated by study designs. In addition, 29 plots revealed new clustering not related to batch information or biological group by study design, suggesting significant sample-to-sample heterogeneity in bulk transcriptome analysis. We identified the relationship of new clustering structures with known sample features for 9 plots. The clustering structures of the rest of 20 plots could result from hidden batch effect or biological features not reported by publications, thus referred to as new clustering with hidden features (**Figure 1b**).

**Figure 1.**
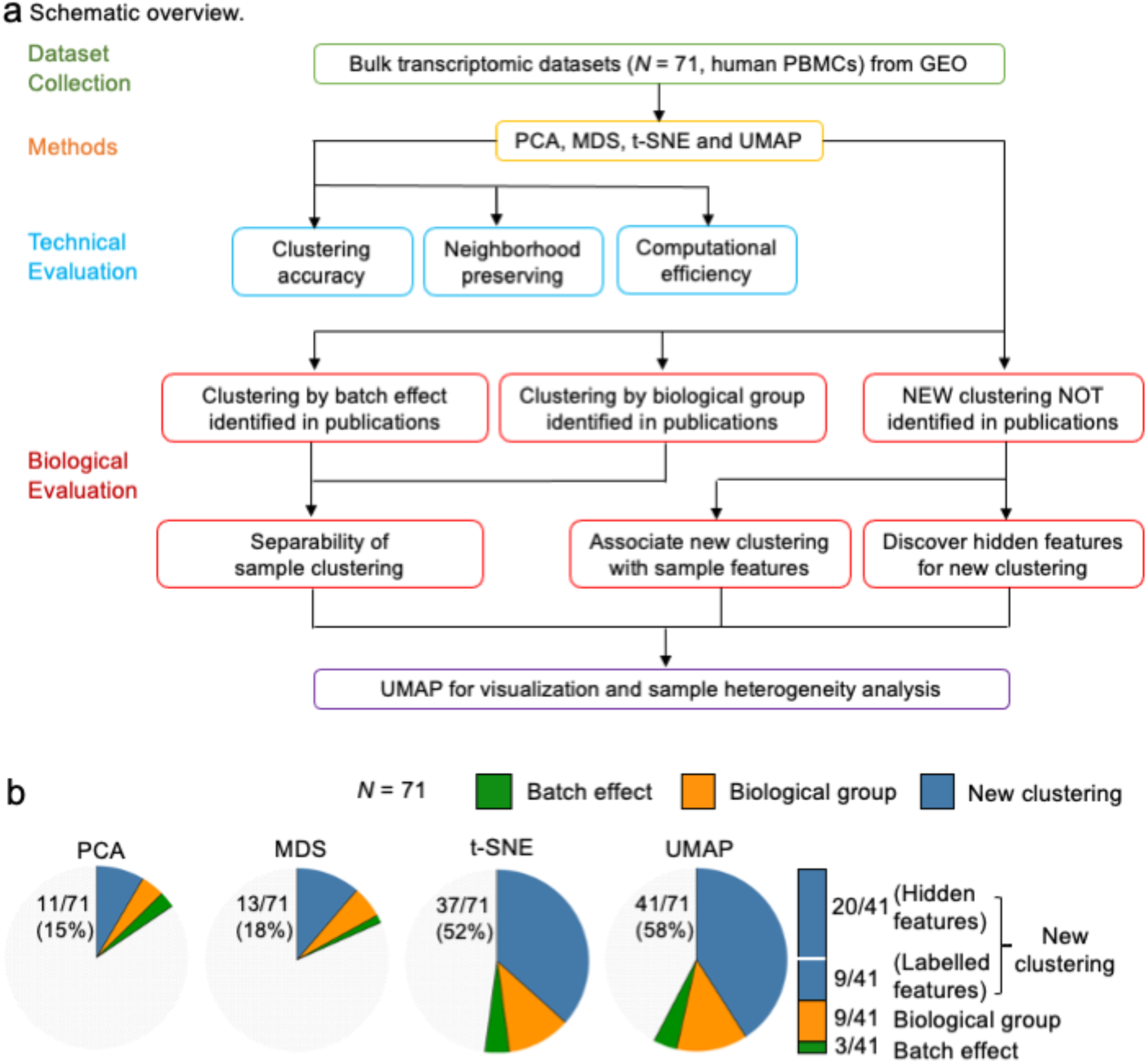
Evaluation overview for four dimensionality reduction methods. (a) Schematic overview of the evaluation. Bulk transcriptomic datasets were collected from GEO database, followed by applying four methods to the datasets for visualization. The methods were evaluated in both technical and biological aspects. Finally, we presented the recommendation on UMAP for visualization. (b) Pie chart showing the percentage of datasets by biological explanations for all revealed clustering structures. By associating features identified in publications, clustering structures were divided into three categories: batch effect (coloured green), biological group (coloured orange) and new clustering (coloured blue). Batch effect was the cluster associated with batch effects. Biological group was related to experimental design like control and treatment groups, while new clustering was the clusters related to other predefined features like gender. New clustering was further divided into new clustering with sample features and new clustering with hidden features by considering available feature information.

With clustering structures generated by PCA, MDS, t-SNE and UMAP, we could evaluate individual methods’ performance for clustering accuracy, local information preservation, and computational efficiency. For datasets with clustering structure by batch effect or biological group, we would then compare the separability of each method in detecting distinct groups. For new clustering structures, we would investigate the relationships of clustering structures with sample features. Based on these quantitative and qualitative assessments, we could provide the recommendation of the best performing method for dimensionality reduction in sizeable bulk transcriptome analysis (**Figure 1a**).

### Comparison of dimensionality reduction methods by quantitative analysis

#### Clustering accuracy

The foremost objective of dimensionality reduction for bulk transcriptomic analysis is to conspicuously distinguish clustering structures of samples which associate biological meaning. We applied five clustering algorithms (k-means, hierarchical clustering, spectral clustering, Gaussian mixture model and hdbscan, with details of five algorithms in **Table S3**) to low-dimensional spaces projected by dimensionality reduction methods and compared the clustering accuracy.

The five clustering algorithms were performed on the embedding two-dimensional coordinates of 22 datasets which have available label information for groups (labelled in **Table S1**). To assess clustering accuracy of dimensionality reduction methods, we then computed Normalized Mutual Information (NMI) [38] and Adjusted Rand Index (ARI) [39] for comparing the true group labels and inferred group labels obtained by clustering algorithms based on the low-dimensional components, and the lager score indicates better clustering accuracy. UMAP was scored the highest for both NMI and ARI, no matter what clustering algorithm used, achieving the best accuracy for clustering (**Figure 2a** and **S1**). t-SNE was scored slightly lower than UMAP but well outperformed MDS and PCA (**Figure 2a** and **S1**).

**Figure 2.**
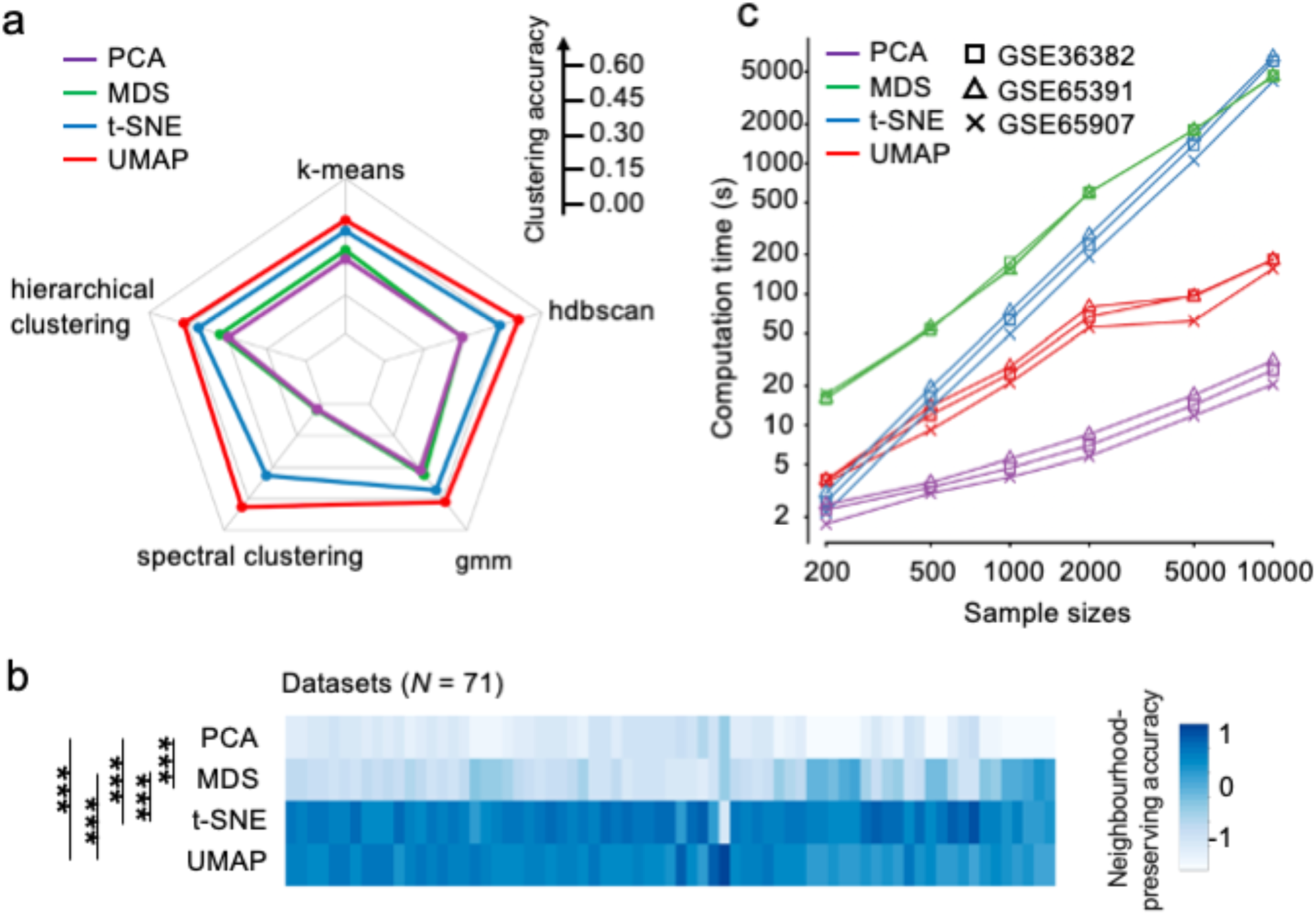
Quantitative analysis of four dimensionality reduction methods. (a) Radar plot of clustering accuracy (average NMI score) comparison using five clustering methods on 22 datasets with cluster labels. The input was the embedded two-dimensional coordinates of each dimensionality reduction methods. Larger scale denotes better clustering accuracy. (b) Heatmap for evaluating neighbourhood preserving of each method on 71 datasets. The number of neighbours is set as 15. The darker the colour is, the better the local information is retained. One-way ANOVA shows significant difference among the four methods (*** *p*<0.001). R function *heatmap* in R package *stats* was used for Figure 2b. (c) Running time evaluation of four dimensionality reduction methods with varying sample sizes. Log-transforming of the time was applied. Different sizes of data were generated by sampling with replacement from three largest datasets respectively.

#### Neighbourhood preserving

We then evaluated the performance of different dimensionality reduction methods in retaining local information from original datasets, which was assessed by comparing the fidelity of local neighbourhood structures between the reduced low-dimensional space and the original space using a Jaccard index (details in ‘Methods’) [40]. The Jaccard indexes were computed for 15 neighbours (**Figure 2b**) and 30 neighbours (**Figure S2**), respectively. PCA exhibited the worst performance in preserving neighbourhood information (averaged 0.19 ± 0.067), followed by MDS (averaged 0.26 ± 0.114). The performance of UMAP (averaged 0.35 ± 0.091) appeared comparable to that of t-SNE (averaged 0.36 ± 0.095), and both were better than PCA and MDS. Pairwise t-test was performed between every two methods (**Figure 2b**), and statistically significant differences were detected between group means by one-way ANOVA (F(3, 280) = 57.88, p < 0.001). This was conceivable since UMAP and t-SNE are designed to utilise local information for dimensionality reduction.

#### Computational efficiency

We next measured the execution time of each dimensionality reduction method on data with sample size ranging from 200 to 10,000. The varied scales of data were generated by randomly sampling with replacement from the three largest datasets (GSE36382, GSE65391 and GSE65907). As shown in **Figure 2c**, the variability of consumed time among different datasets was negligible. PCA performed consistently faster than the other three methods while MDS ran slowest (**Figure 2c**). For 200 and 500 samples, consumed time was similar between t-SNE and UMAP but UMAP gained an advantage for data with larger sample sizes. For processing a data with 10,000 samples, UMAP (∼3 minutes) was more than 25-time faster than t-SNE (∼ 1.5 hours), although still slower than PCA (∼20 seconds) (**Figure 2c**). PCA and UMAP appeared more time-efficient than MDS and t-SNE for computing large-sized data.

Technically, UMAP not only identified more clustering structure in 71 datasets of bulk transcriptome analysis (**Figure 1b**), but was also superior to the other three methods for the overall performance by assessing the three quantitative criteria. We next compared four dimensionality reduction methods for uncovering biological meaning.

### Comparison of dimensionality reduction methods by qualitative analysis

#### Identification of batch effects

Batch effects are common in many types of high-throughput sequencing experiments, which are systematic technical variations introduced by processing samples in different batches [6, 41]. As for high-throughput sequencing experiments, it is essential to remove unwanted variations in the transcriptomic analysis by normalisation [42, 43] to avoid biased analysis and distorted results [6]. The first step is to identify batch effects among samples. PCA is the most used tool, such as by The Cancer Genome Atlas (TCGA) project [21]. It generates the clustering structure of samples in two-dimensional embedding space to facilitate the visualisation for batch information. Among the 41 datasets with explicit clustering structures, three datasets showed clustering structures related to batch effects reported by publications (**Figure 1b**). Each dimensionality reduction method was used to visualise batch effects for the three datasets (one in **Figure 3a** and two in **S3**). UMAP and t-SNE showed better segregation between samples from different batches. To assess the ability of each method to separate batch effects in two-dimensional embeddings, we trained random forests to predict batch effects from sample points in embedding space and calculated the prediction accuracy on held-out data (details in ‘Method’). Consistent with the visualisation, UMAP and t-SNE performed better than MDS and PCA, leading to random-forest accuracies around 90% (**Figure 3b**).

**Figure 3.**
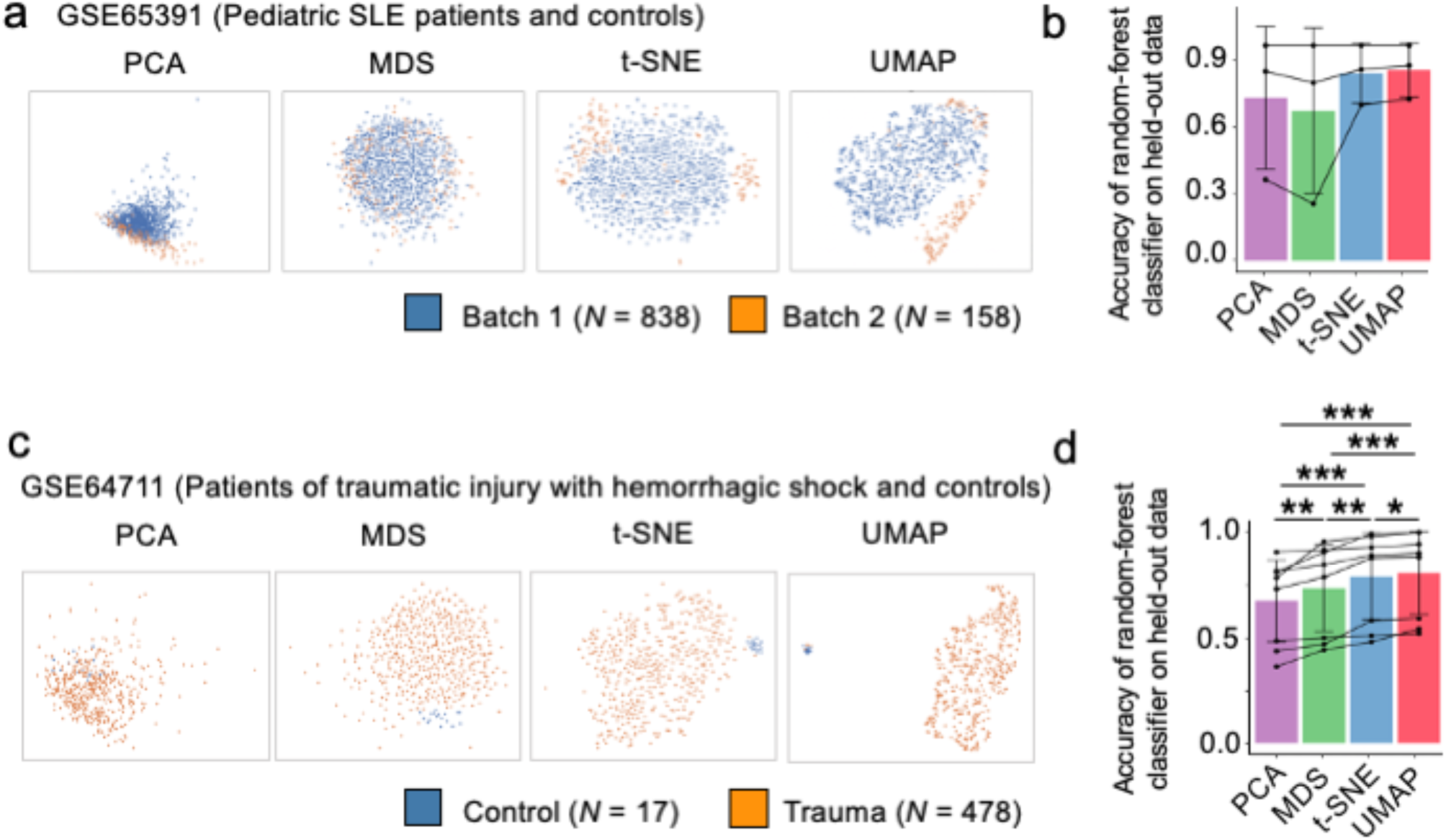
Biological explanation of clustering by batch effects and biological group. (a) Visualization of dataset GSE65391 showing the batch effects (coloured by blue and orange) in two-dimensional space by dimensionality reduction methods. (b) Visualization of dataset GSE55447 illustrating biological group by dimensionality reduction methods. Control group is labelled in blue and trauma group is in orange. (c), (d) Classification accuracy on held-out data of random-forest classifiers predicting cluster labels taking embedded coordinates as input. (c) is for batch effects, while (d) for biological groups. The average score across datasets is shown, with vertical bars representing s.d.; paired t-test was conducted on pairwise methods (∗ p < 0.05, ∗∗ p < 0.01, ∗∗∗ p < 0.001)

#### Validation of biological groups by experimental design

One major purpose of bulk transcriptome analysis is for the DGE analysis between biological groups defined by experimental design. Visualising the segregation of samples from groups with distinct biological features by dimensionality reduction is often applied to the validation of group-to-group distinction. Among the 41 datasets with explicit clustering structures, 9 datasets showed clustering structures related to biological groups by experimental designs (**Figure 1b**). We compared four dimensionality reduction methods in visualizing biological group and found that UMAP and t-SNE outperformed MDS or PCA in visually separating biological groups in 9 datasets (one in **Figure 3c** and eight in **S4**). To measure the separability of each method in group validation, we again deployed random forests to train embedding data with group features as labels and computed the prediction accuracy on held-out data (details in ‘Method’). UMAP achieved the best accuracy (> 80%) than t-SNE (*p* < 0.05), MDS (*p* < 0.001) and PCA (*p* < 0.001) in separating biological groups (**Figure 3d**).

#### Uncovering new associations between clustering structures and sample features

Only 12 out of 41 datasets showed clustering structures explained by batch effects or biological groups (**Figure 1b**). The appearance of new clustering structures in 29 plots demonstrated significant heterogeneity existing in bulk transcriptome profiling, which could be efficiently revealed by UMAP. We next investigated the causes underlying new clustering structures. The clustering structures in 9 datasets were found associated with certain sample features reported by publications (**Figure S5**). These features were not used for the classification of sample groups in experimental designs, suggesting certain biological features with major impacts on sample heterogeneity were not included in experimental designs or data analyses. A good case was the dataset GSE71220, which was designed to determine the impact of cigarette smoking (former v.s. current smoker) on gene expression in peripheral blood of patients with chronic obstructive pulmonary diseases (COPD) [44]. Dimensionality reduction methods of UMAP and t-SNE generated plots showing clustering structures (right part in **Figure 4a**). However, such clustering was not associated with smoking status (**Figure 4b**). We applied other sample features including age and disease status to the two-dimensional plots. Surprisingly, the sample feature of gender demonstrated clear association with clusters in the plots generated by UMAP and, to less extent, t-SNE (**Figure 4c**). In the UMAP plot, one cluster was highly enriched of females (in orange colour) and another cluster was highly enriched by males (in blue colour), with the third cluster showing the pattern of a mixture (**Figure 4c**). By deploying spectral clustering (details in ‘Methods’), samples were divided into three clusters with distinct gender composition: C1-97% females, C2-93% males, and C3-mixed (**Figure 4d, e**). This indicated that the transcriptomes of samples in this study were highly influenced by gender difference. Indeed, the heatmap of the top 100 differentially expressed genes demonstrated that the clustering of samples was strongly associated with gender (**Figure 4f**). Therefore, the heterogeneity uncovered by the dimensionality reduction using UMAP indicated that the gender difference should have been critically treated as a latent variable in downstream transcriptomic analysis.

**Figure 4.**
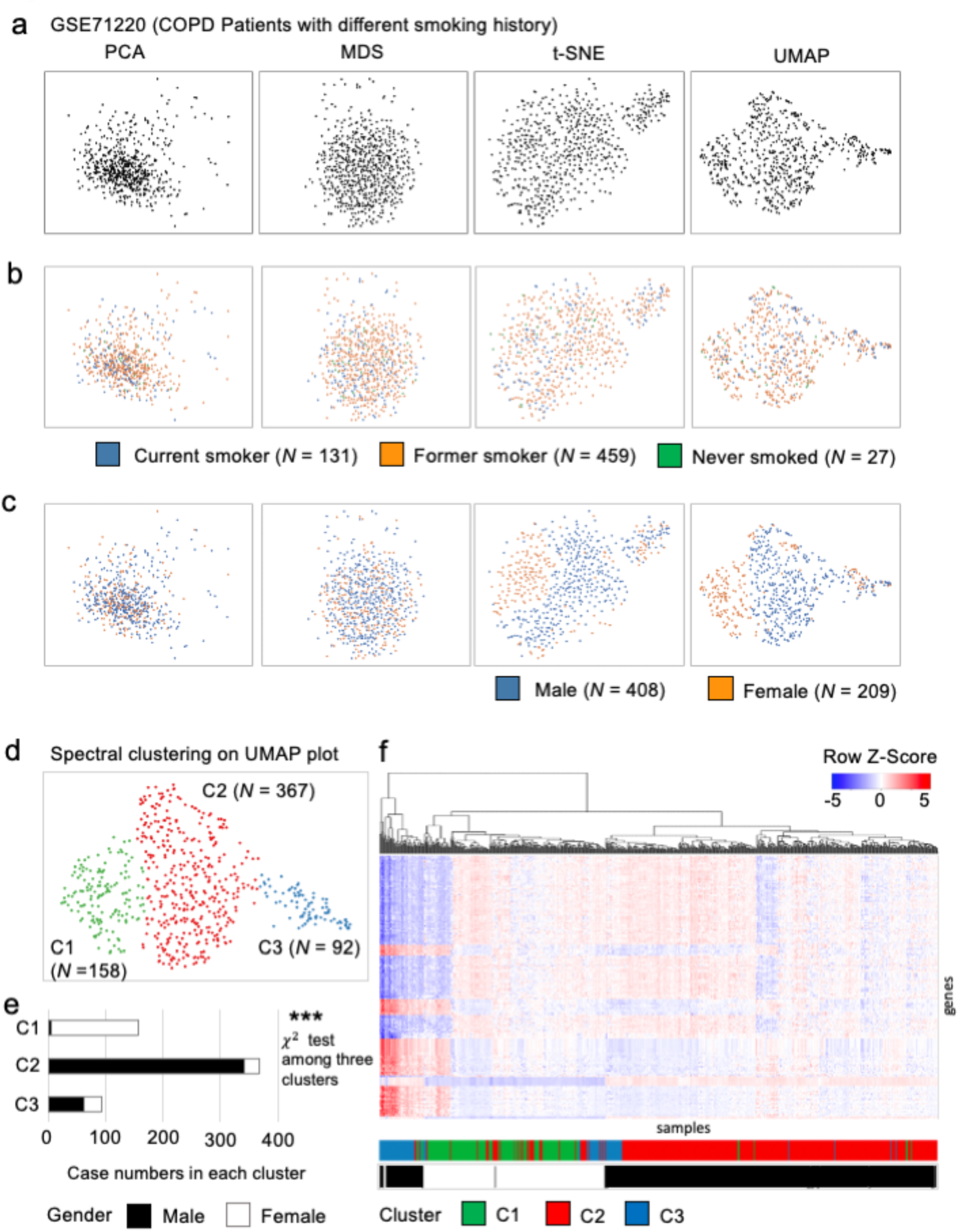
New clustering interpreted by available sample features. (a)-(c) Visualization of dataset GSE71220 in two-dimensional space by assigning no feature (a), group labels (b), gender (c). (d) Spectral clustering on two-dimensional embedded coordinates into three clusters: C1, C2, C3. (e) Gender proportion among three clusters by χ^2^ test showing a significant difference (*** *p*<0.001). Male and female are coloured by black and white respectively. (f) Heatmap of top-100 differentially expressed genes with three clusters C1, C2, C3 and two gender groups male and female. R function *heatmap*.*2* in R package *gplots* was used for Figure 4f.

#### Discovering new associations between clustering structures and hidden features

By dimensionality reduction using UMAP, 41 datasets showed clustering structures in two-dimensional embedding spaces in which the associations with batch effects, biological groups by experimental designs or specific sample features reported by publications were identified in 21 datasets (**Figure 1b**). For the rest 20 datasets, clustering structures might derive from obscure heterogeneity of samples, biologically or technically (**Figure S6**). We made efforts to explore the biological meanings of clustering structures of these datasets and herein present the dataset GSE121239 as an example to support the notion that new clustering structures generated by UMAP can reinforce sample heterogeneity analysis of bulk transcriptome data to reveal important biological meaning.

Dataset GSE121239 originated from the study of systemic lupus erythematosus (SLE) which is the prototype of systemic autoimmune diseases with highly diverse manifestations in multiple tissues and organs, such as skin, kidney and lung [45]. As a chronic disease, SLE patients often experience unpredictable occurrence of disease flares [46]. In order to identify the heterogeneity of SLE patients and stratify patient groups of disease activity progression, the dataset GSE121239 collected longitudinal transcriptome profiles of 65 SLE patients with more than three clinical visits and 20 healthy individuals as controls [47]. Data collected at each visit contributed to one sample in the dataset. Dimensionality reductions plot by UMAP and t-SNE, but not PCA or MDS, demonstrated clearly separated clusters for SLE patients (in orange colour) and healthy controls (in blue colour) (**Figure 5a, b**). In the UMAP plot, we noticed more than one cluster for patient samples (**Figure 5c**). To understand the biological meaning of clusters representing subgroups of SLE patients, we examined feature information of patients reported by the publication including gender and patient ID but found no direct association with the clustering structure of patient subgroups. Since the samples of patients were collected longitudinally from multiple clinical visits, we set samples collected at the first clinical visit as day 1 then labelled subsequently collected samples from the same patient with the period between two visits. The resulted contour plot showed samples in the chronological order (**Figure 5d**). Importantly, the gradient from light to dark orange spreads from the middle of the plot to two sides, indicating the clustering structure generated by UMAP was associated with the timing evolution of clinical visits. For example, the bottom-right cluster in **Figure 5c** represents samples collected from a subgroup of patients at their late clinical visits, indicated by dark orange in **Figure 5d**. This intriguing discovery suggested that new clustering structures revealed by UMAP could facilitate the exploration of samples’ hidden features.

**Figure 5.**
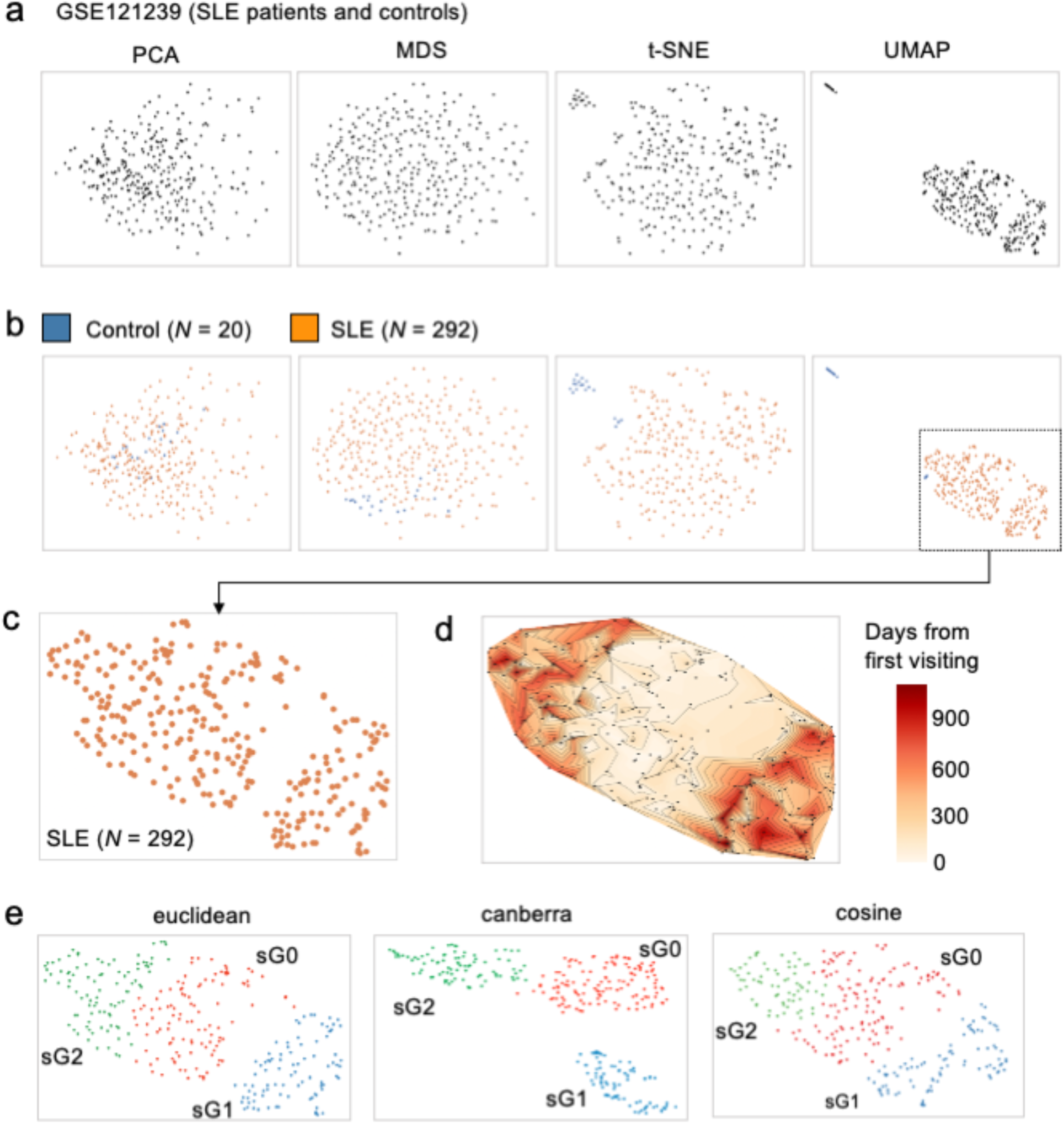
Discovering new associations between clustering structures and hidden features. (a), (b) Visualization of dataset GSE121239 in two-dimensional space by assigning no feature (a), group labels (b). (c) Patient (SLE) group (coloured orange) showing new clustering structure (sG1, lower right). (d) Contour plot on patient groups by the order of visiting timestamp. Each data point is associated with one visiting timestamp. Data points are coloured by the order of visiting time with light colours for early visits and dark colour for late visits. The code to plot Figure 5d is in Code Availability. (e) Hierarchical clustering of patient group on two-dimensional embedded coordinates by UMAP with distance metric as ‘euclidean’, ‘canberra’, and ‘cosine’, respectively.

To generate UMAP plots, there are several options for metric space, with ‘*euclidean*’ distance as default [18]. We tested ‘*euclidean*’ and another two representative metrics ‘*canberra*’ and ‘*cosine*’ and observed that the metric ‘*canberra*’ led to more explicit clustering on UMAP projection, with patients’ samples clustered into three subgroups: sG0, sG1, sG2 (**Figure 5e**).

According to the timing evolution (**Figure 5d**), samples of sG0 were collected earlier while samples of sG1 or sG2 were collected later. The clear separation of late collected samples into two clusters of sG1 and sG2 suggested a biological divergence. To interpret the biological difference between sG1 and sG2, we applied gene set enrichment analysis (GSEA) using R package EGSEA [48], resulting in the top 20 differentially regulated molecular pathways between sG1 v.s. sG0 and sG2 v.s. sG0 (**Figure 6b, S7**). Comparing to sG0, sG1 and sG2 were common in 6 upregulated pathways (in red colour) and 2 down-regulated pathways (in blue colour). However, 7 upregulated and 5 downregulated pathways in sG1 showed opposite trends in sG2, suggesting the biological distinction between them.

Given longitudinal sampling of individual patients, we next investigated the visit trajectories of individual patients. Connection of samples from each patient demonstrated that most patients (*N* = 47/65) showed one-directional trajectories from sG0 to sG1 or sG0 to sG2 (**Figure 6c**), in agreement with the timing evolution of patients’ sample (**Figure 5d**). When initially admitted to the clinic to take samples (visit 1, **Figure 6d**), patients with distinct trajectories had comparable disease activities (SLE disease activity index (SLEDAI), mean±SD, sG0 to sG1: 2.6±2.71; sG0 to sG2: 2.6±2.85). Widely used in clinical practice and research, SLEDAI is a global index that was developed as a clinical index for the assessment of lupus disease activity and larger SLEDAI indicates worse disease conditions [5]. Importantly, we noticed that the average SLEDAI at the following visits increased for patients with the trajectory from sG0 to sG1 (in blue colour, **Figure 6d**), indicating the disease deterioration of these patients, whereas the average SLEDAI at the following visits decreased for patients with the trajectory from sG0 to sG2 (in green colour, **Figure 6d**), indicating the disease improvement of these patients. The opposite disease progression between two trajectories was also supported by GSEA, which showed the key pathogenic pathways for SLE including apoptosis [49], type I interferon [50] and type II interferon [51] were increased in sG1 but decreased in sG2 (**Figure 6b**). Taken together, the deep exploration of the biological and clinical meaning of the new clustering structure of dataset GSE121239 revealed by UMAP supports the future application of dimensionality reduction methods such as UMAP to reinforce sample heterogeneity analysis of bulk transcriptome data.

**Figure 6.**
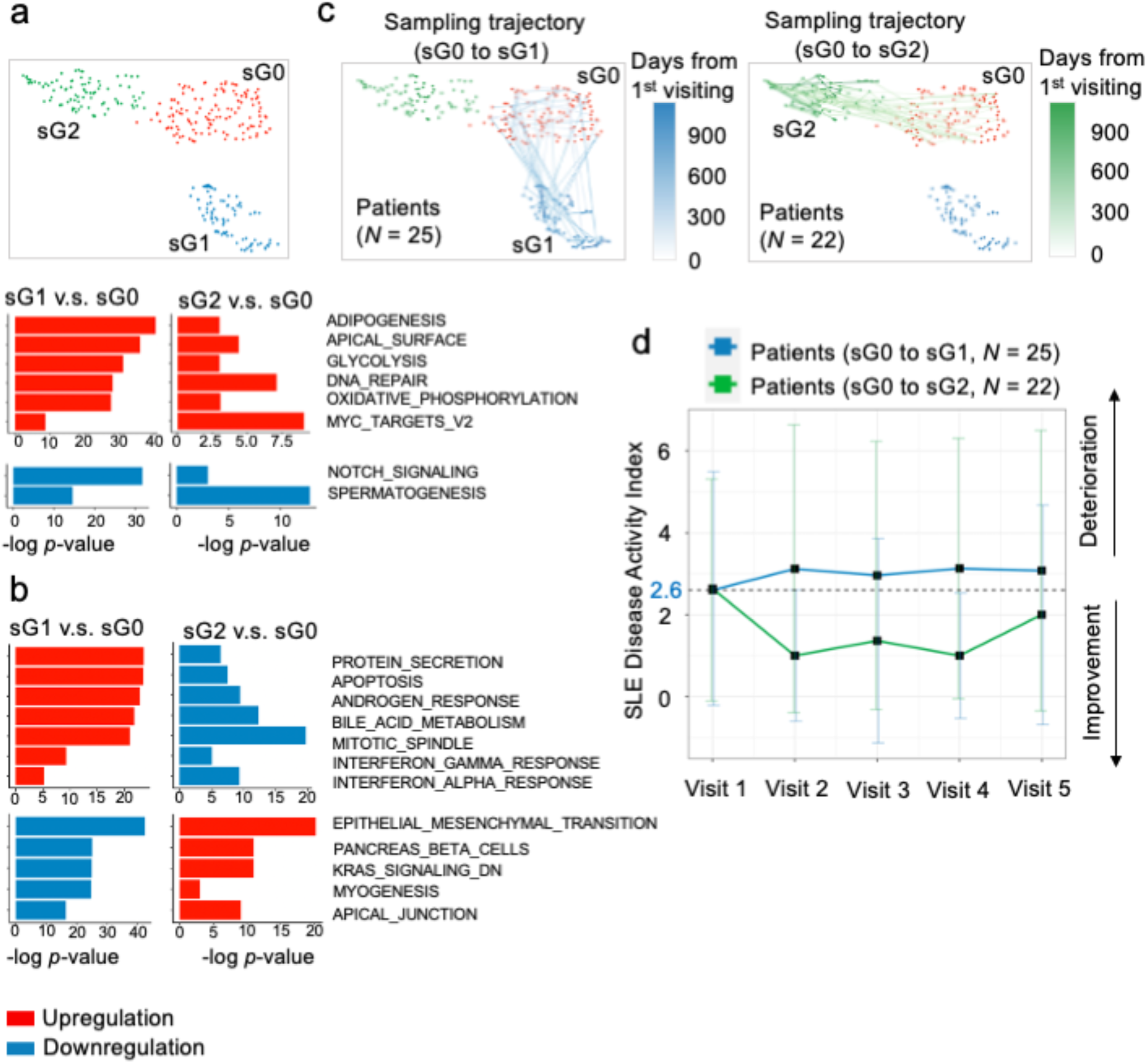
UMAP revealed clustering structure explained by clinical traits. (a) Hierarchical clustering of patient groups on two-dimensional embedded coordinates by UMAP with metric as ‘canberra’. (b) Histogram illustrating gene set enrichment analysis between sG1 v.s. sG0 and sG2 v.s. sG0 with top 20 differentially regulated molecular pathways (negative logarithm of the *p*-value (base 10)). Colour red denotes upregulation and blue for downregulation. The top two rows are the same direction of regulation, and the bottom two rows are in the opposite direction. (c) Visiting trajectories of each patient on UMAP plot with metric = ‘canberra’. Each path connected data points corresponding to one patient with several visits. Data points in pathes were connected by visiting timestamp. The light colour denotes early visit and dark colour for late visits. The paths were mainly divided into two patterns: from sG0 to sG1, from sG0 to sG2. (d) Line chart of average SLEDAI changing along with visits between sG0 to sG1 and sG0 to sG1. Both started with average SLEDAI around 2.6; from sG0 to sG1 (coloured by blue) the average SLEDAI increased, while from sG0 to sG2 (coloured by green) the average SLEDAI decreased.

### Recommendation

Although PCA is often used in identifying sample-to-sample heterogeneity in bulk transcriptome analysis, our study demonstrated that the nonlinear dimensionality reduction method UMAP improved the identification, visualisation and interpretation of clustering structures in sizeable datasets. The analysis of the dataset GSE121239 suggested that the choice of the parameter ‘*metric*’ in UMAP could affect the visualisation of clustering structures of UMAP plots (**Figure 6a**). We then thoroughly evaluate ‘*euclidean*’, ‘*canberra*’ and ‘*cosine*’ metrics of UMAP in all 71 bulk transcriptomic datasets, which respectively revealed clustering structures in 41, 44 and 42 datasets and had 39 datasets in common (**Figure 7a**). Without any ‘*metric*’ showing a clear advantage, we recommend trying the three representative metrics for UMAP in visualising the bulk transcriptomic data and being integrated into the pipeline for bulk transcriptomic analysis (**Figure 7b**). The analysis starts with transcript counts as the input, followed by applying UMAP to visualise potential clustering structures. If no clustering structure is detected, DGE analysis can be performed. With clustering structures that may correspond to known or unknown batch effects, the first consideration is to identify and remove batch effects. The clustering structure should next be tested for the association with biological groups assigned by experimental design. The explicit association of the clustering structure with biological groups can ensure robust DGE analysis among different biological groups. If the clustering structure is related to specific sample features rather than biological groups, that feature should be treated as latent covariates in DGE analysis. On the other hand, the clustering structure might reveal new biological subgroups or hidden factor to be analysed separately for DGE analysis.

**Figure 7.**
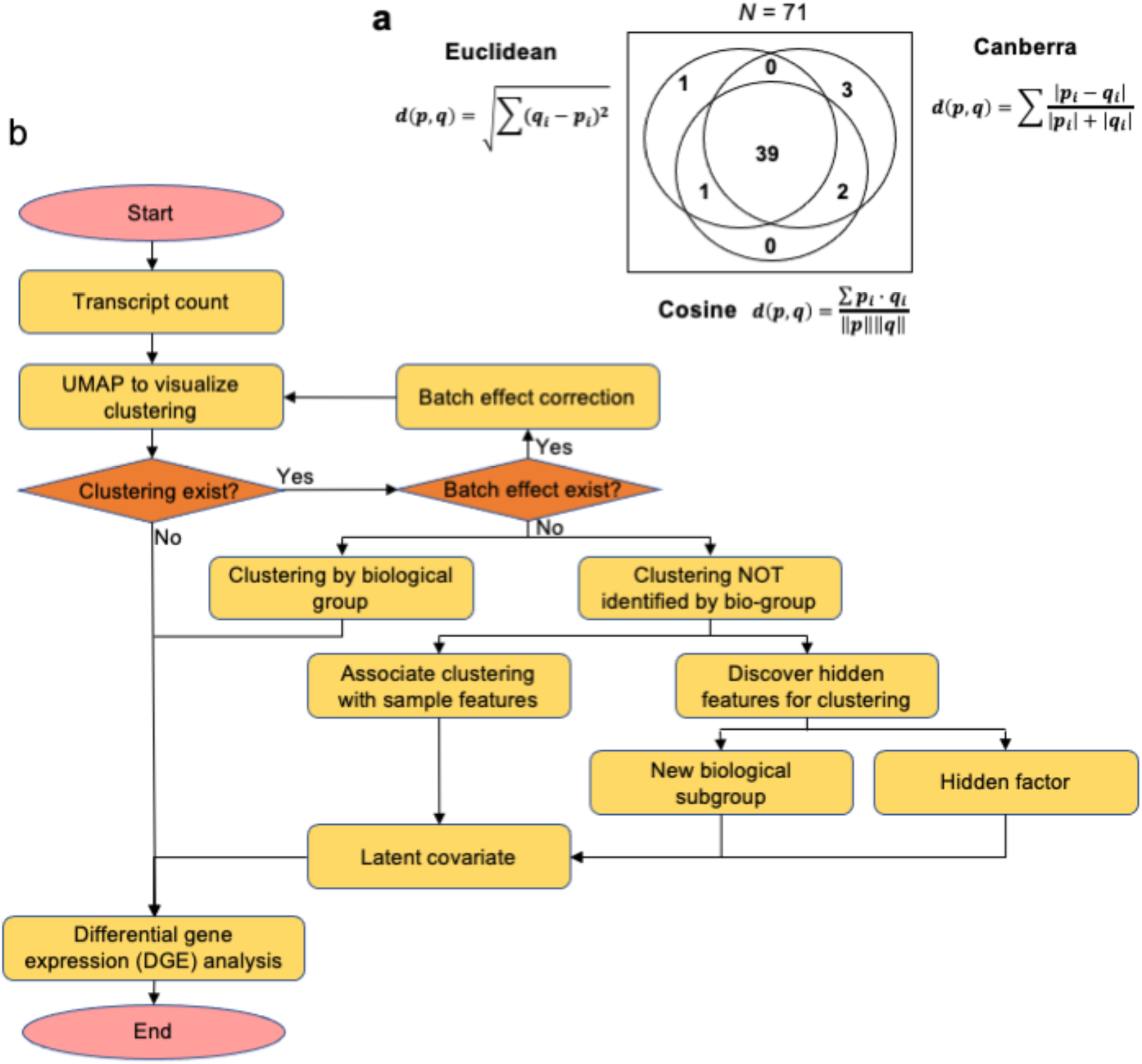
Recommendations for UMAP processing bulk transcriptome datasets. (a) Venn diagram illustrating the overlap in the number of datasets having clustering structure by the UMAP plot under three different ‘*metric*’ parameters: ‘*euclidean*’, ‘*canberra*’, and ‘*cosine*’. (b) The recommendation pipeline for applying UMAP to bulk transcriptome analysis.

## Discussion

Sample heterogeneity in bulk transcriptomic data reflects both biological and technical variation among samples. It is crucial to detect among-sample heterogeneity before DGE analysis for bulk transcriptomic data so that appropriate analytic methods can be subsequently used to correct batch effects, remove outliers and distinguish subgroups. Sample heterogeneity analysis by dimensionality reduction should consider both local and global information of datasets to congregate similar samples and distinguish different samples. PCA is the current mainstream tool of dimensionality reduction to visualise and detect among-sample heterogeneity, adopted by widely used analytic packages limma and edgeR [11, 12]. PCA produces linear combinations of the original variables to generate the principal components [52], and visualisation is generated by projecting the original data to the first two principal components, thus PCA plot linearly shows global distance among data points. Similarly, MDS method places each data point into two-dimensional space such that the between-point distances are preserved according to the pairwise distance of original data points [53]. Both PCA and MDS focus more on maintaining global information, which can fail to compactly cluster similar data points and face a major challenge with the rapid increase in sample sizes of bulk transcriptomic profiling datasets.

On the other hand, t-SNE and UMAP model the pairwise distance by adopting the concept from k-nearest neighbour (kNN) graph [17, 18] whereby two points are connected by an edge if their distance is among the k-th smallest distances compared to distances to other points [54]. For dimensionality reduction by t-SNE or UMAP, all pairs of two points have edge weights indicating the probability for them being connected (connection probability). If the distance between two points is among the k-th smallest distances compared to distances to other points, the connection probability between these two points is high. If the distance between two points is much greater than the k-th smallest distance, the connection probability between these two points is low [17, 18]. Therefore, t-SNE and UMAP can efficiently preserve local distance information and cluster similar sample points. For large sample size in dataset resulting in the quadratic increase of pairwise comparisons, t-SNE and UMAP not only retain pairwise interaction but also focus on local information, thus outperforming PCA and MDS in detecting sample heterogeneity. Compared with t-SNE using random initialisation and KL-divergence object function, UMAP utilises Laplacian Eigenmaps initialisation and cross-entropy object function [18, 55] which contribute to the global structure preservation. This might explain the overall better performance of UMAP than t-SNE. We tested three presentative parameters for the distance ‘*metric*’ of UMAP – ‘*euclidean*’, ‘*canberra*’ and ‘*cosine*’ and found consistent outcomes with only minor variation (**Figure 7a**).

Among 71 bulk transcriptome profiling datasets with > 100 samples tested in this study, UMAP and t-SNE clearly outperformed PCA and MDS in identifying clusters associated with batch effects and biological groups pre-defined in study designs. It should be noted that, within 41 of 71 datasets that UMAP identified clustering structures, new fine-scale clustering structures were revealed and accounted for more than half (29 out of 41) (**Figure 1**). The important question is whether the new clustering structures discovered by UMAP represent biological significance. This question was then addressed in case studies of datasets with new clustering structures. One case is the study that was initially designed to investigate how smoking influence blood gene expression of patients with COPD and utilised bulk transcriptomic profiling and DGE analysis (GSE71220 [44]). Intriguingly, the PCA plot showed no clustering structure while the UMAP plot revealed new clustering structures, which was related to gender rather than smoking status (**Figure 4**). This information discovered by dimensionality reduction using UMAP suggests the gender feature should be treated as an important latent covariate in DGE analysis. Another example is the study that was designed to stratify patients with SLE, a highly complex autoimmune disease with heterogeneous clinical presentation, according to longitudinal disease activity and blood gene expression (GSE121239 [47]). This study calculated a gene-by-patient correlation matrix computing a stringent Pearson correlation coefficient between gene expression data and SLEDAI scores across each patient’s visits and then selected genes with the highest absolute correlation values by rank-sum method [47]. Instead of this multiple-step process, dimension reduction by UMAP revealed the separation of samples by visit timestamp (**Figure 5**), which enabled to identify two groups of patients with opposite changes of longitudinal disease activity (**Figure 6**). These results thus validate the application of UMAP in dimensionality reduction in stratifying SLE patients. Using several datasets as examples, we demonstrated that the new clustering structures were associated with certain sample features and enabled to uncover unappreciated sample subgroups with specific biological and clinical features.

In analysing 71 datasets, we demonstrated that UMAP was able to visualise the among-sample heterogeneity in two-dimensional space. Based on the low-dimensional embedding space of UMAP, clustering methods were deployed to define clusters of the data points (**Figure 4d** and **Figure 5e**). The biological significance of resulting clusters was validated by subsequent exploration and evaluation (**Figure 4** and **Figure 6**). For scRNA-seq data, clustering algorithm is generally applied on low-dimensional space, for example in the commonly used scRNA-seq package *Seurat* [57], a graph-based clustering algorithm to low dimensional space by PCA projection. The rationale of applying clustering method to low-dimensional projected space mainly arises from the curse of dimensionality [56]. When computing distance (e.g., Euclidean distance) in high-dimensional data, the difference in the distances between different pairs of samples becomes less precise, which hinders discriminating near and far points. Thus, applying clustering methods to low-dimensional embedding space is better to define clusters of data points. Therefore, we suggest that UMAP can be applied as a pre-processing step before generating clusters from bulk transcriptomic datasets.

Although UMAP has shown significant advantages in detecting among-sample heterogeneity. PCA has a unique property not present by other methods. PCA compresses the data by top-ranked principal components and computes the PCA score for each sample. Therefore, it can calculate the variable weight corresponding to new coordinate system (PCA loadings), which explains the contribution of each variable to sample points. In contrast, the nonlinear methods, including MDS, t-SNE and UMAP, do not involve the variable weight such that dimensionality reduction embedding cannot be immediately explainable by variable weight. This might represent an area for the future improvement of UMAP or methods of similar kind.

Though commonly used for scRNA-seq, UMAP has been repurposed in large scale genotype datasets to explore the fine structure and visualise genetic interactions [59, 60]. Based on the quantitative and qualitative results of the comparison among dimensionality reduction methods, we highly recommend UMAP as the visualisation tool in the pipeline for bulk transcriptomic profiling and DGE analysis. It can particularly reinforce sample heterogeneity analysis for datasets with large sample sizes.

## Methods

### Datasets

The total RNA datasets were collected from the Gene Expression Omnibus (GEO) database with query conditions set as follows: the dataset type was expression profiling by array or by high throughput sequencing; the number of samples ranged from 100 to 10,000; organism was homo sapiens; the publication date was from 2015/01/01 to 2020/03/01; sample source was PBMC or whole blood. Applying the query to the GEO database, we gained 214 results. We further manually removed the datasets in which each group owned less than 100 samples, resulting in 71 datasets.

### Clustering accuracy (NMI, ARI)

For clustering accuracy analysis, we applied five clustering methods to the embedded low-dimensional space by dimensionality reduction methods. The clustering methods included k-means clustering (Python function *KMeans*), hierarchical clustering (Python function *AgglomerativeClustering*), spectral clustering (Python function *SpectralClustering*), hdbscan (Python function *hdbscan*) and Gaussian mixture model (Python function *GaussianMixture*). In these clustering methods, the number of clusters *k* was set to be the known number of different groups in the data, except for hdbscan which is a density-based clustering algorithm (we set the *min_cluster_size* as 10). We applied the five clustering methods to the embedded space of 26 datasets with available features for groups. The retained partitions inferred using the low-dimensional components were compared to the true clusters. The level of agreement between the clustering partition and the true clusters was measured by two criteria: the Adjusted Rand Index (ARI) [39] and the Normalized Mutual Information (NMI) [38]. Given two partitions *X* = {*X*_1_,…, *X*_*r*_} and *Y* = {*Y*_1_,…, *Y*_*s*_}, the ARI and NMI are defined as:

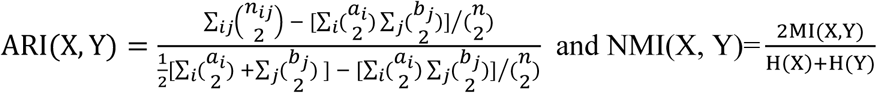

where *n*_*ij*_ = |*X*_*i*_ ∩ *Y* _*j*_| is the number of common data points between *X*_*i*_ and *Y* _*j*_, *a*_*i*_ = ∑_*j*_ *n*_*ij*_, *b* _*j*_= ∑_*i*_ *n* _*ij*_, MI(X, Y) is the mutual information between cluster labels *X* and *Y*, H(X) and H(Y) are the entropy function for cluster labelling. We used Python function *adjusted_rand_score* and *normalized_mutual_info_score* to calculate ARI and NMI, respectively.

### Neighbourhood preserving evaluation

The evaluation of neighbourhood preserving is to assess how the reduced low-dimensional space retains the local information compared with the original high dimensional dataset. For the original space and embedded space, the k-nearest neighbours (kNNs) for each data point were computed respectively (denoted as sets X and Y). The Jaccard index (JI) [40] was used to calculate the neighbourhood similarity between original and embedded space: *JI*=|X∩Y|/|X∪Y| where |⋅| means set cardinality, then the average Jaccard index (AJI) across all data points were computed to measure the neighbourhood preserving.

### Running time

We measured the running time of PCA, MDS, t-SNE and UMAP on a single thread of an Intel Xeon E5-2698 v4 2.20GHz processor. The running time was determined in R using the “elapsed” (wall clock) time measurements, which allows for consistent timing across methods. For total-RNA datasets, the number of samples is moderately large with hundreds of data points. We generated datasets with sample size ranging from 200 to 10000 by random sampling to evaluate the computation efficiency. The data were generated by randomly sampling with replacement from the three largest datasets (GSE36382, GSE65391 and GSE65907).

### Separability of batch effects and biological groups

To evaluate the capability of each dimensionality reduction method in separating the groups by the embeddings, we first assigned batch labels to 3 datasets and biological group labels to 9 datasets. For each dataset, we used Python function *train_test_split* with parameter *test_size = 0*.*3* to divide the dataset into 70% training set and 30% test set. For each algorithm, a random-forest classifier by Python function *RandomForestClassifier* was trained using the group labels as target variable and the embedding’s coordinates as training variables. We then utilized these classifiers to predict cluster identities on the test set and computed the accuracy of these predictions, thus assessing the ability of each method to separate groups.

### Statistical test

We applied two-tailed t-test to compare the performance of dimensionality reduction methods. The frequency difference of categorical variables was examined by χ^2^ test. The p-value less than 0.05 is considered statistically significant. We used R (3.6.3) package *limma* [11, 13] for differential gene expression (DGE) analysis. Top 100 differential expressed genes were chosen to be included in the heatmap among control and experimental groups. We applied gene set enrichment analysis (GSEA) by R package *EGSEA*, where the Molecular Signatures Database (MSigDB) was set as H: hallmark gene sets [61].

### Data availability

The datasets supporting the conclusions of this article are available in Gene Expression Omnibus repository (https://www.ncbi.nlm.nih.gov) with the GEO accession numbers in **Table S1**, including four columns (UMAP, t-SNE, MDS and PCA) showing which feature information explains the clustering structure of each dataset.

### Code availability

All scripts used for dimensionality reduction and clustering are available through Github https://github.com/yuImmuGroup/umap_on_bulk_transcriptomic_analysis; differential gene expression and gene set enrichment analysis are available in https://github.com/yuImmuGroup/transcriptomic_analysis_DGE_and_GSEA.

## References

1. Heller, M.J., DNA microarray technology: devices, systems, and applications. Annual review of biomedical engineering, 2002. 4(1): p. 129–153.

2. Lim, E., et al., Aberrant luminal progenitors as the candidate target population for basal tumor development in BRCA1 mutation carriers. Nature medicine, 2009. 15(8): p. 907–913.

3. Wang, Z., M. Gerstein, and M. Snyder, RNA-Seq: a revolutionary tool for transcriptomics. Nature reviews genetics, 2009. 10(1): p. 57–63.

4. Wu, D., et al., Gene-expression data integration to squamous cell lung cancer subtypes reveals drug sensitivity. British journal of cancer, 2013. 109(6): p. 1599–1608.

5. Bombardier, C., et al., Derivation of the SLEDAI. A disease activity index for lupus patients. Arthritis & Rheumatism: Official Journal of the American College of Rheumatology, 1992. 35(6): p. 630–640.

6. Leek, J.T., et al., Tackling the widespread and critical impact of batch effects in high-throughput data. Nature Reviews Genetics, 2010. 11(10): p. 733–739.

7. MDAndersonCancerCenter, U.A. TCGA Batch Effects Viewer. 2020 [cited 2020 1st October]; Available from: https://bioinformatics.mdanderson.org/public-software/tcga-batch-effects/.

8. Wold, S., K. Esbensen, and P. Geladi, Principal component analysis. Chemometrics and intelligent laboratory systems, 1987. 2(1-3): p. 37–52.

9. Torgerson, W.S., Multidimensional scaling: I. Theory and method. Psychometrika, 1952. 17(4): p. 401–419.

10. Love, M.I., W. Huber, and S. Anders, Moderated estimation of fold change and dispersion for RNA-seq data with DESeq2. Genome biology, 2014. 15(12): p. 550.

11. Ritchie, M.E., et al., limma powers differential expression analyses for RNA-sequencing and microarray studies. Nucleic acids research, 2015. 43(7): p. e47–e47.

12. Robinson, M.D., D.J. McCarthy, and G.K. Smyth, edgeR: a Bioconductor package for differential expression analysis of digital gene expression data. Bioinformatics, 2010. 26(1): p. 139–140.

13. Law, C.W., et al., RNA-seq analysis is easy as 1-2-3 with limma, Glimma and edgeR. F1000Research, 2016. 5.

14. Tang, F., et al., mRNA-Seq whole-transcriptome analysis of a single cell. Nature methods, 2009. 6(5): p. 377–382.

15. Hwang, B., J.H. Lee, and D. Bang, Single-cell RNA sequencing technologies and bioinformatics pipelines. Experimental & molecular medicine, 2018. 50(8): p. 1–14.

16. Van Buren, E., et al., TWO-SIGMA: a novel TWO-component SInGle cell Model-based Association method for single-cell RNA-seq data. bioRxiv, 2020: p. 709238.

17. Maaten, L.v.d. and G. Hinton, Visualizing data using t-SNE. Journal of machine learning research, 2008. 9(Nov): p. 2579–2605.

18. McInnes, L., J. Healy, and J. Melville, Umap: Uniform manifold approximation and projection for dimension reduction. arXiv preprint arXiv:1802.03426, 2018.

19. Becht, E., et al., Dimensionality reduction for visualizing single-cell data using UMAP. Nature biotechnology, 2019. 37(1): p. 38–44.

20. Consortium, G., The Genotype-Tissue Expression (GTEx) pilot analysis: Multitissue gene regulation in humans. Science, 2015. 348(6235): p. 648–660.

21. Tomczak, K., P. Czerwińska, and M. Wiznerowicz, The Cancer Genome Atlas (TCGA): an immeasurable source of knowledge. Contemporary oncology, 2015. 19(1A): p. A68.

22. Lou, J., et al., Rad18 mediates specific mutational signatures and shapes the genomic landscape of carcinogen-induced tumors in vivo. NAR Cancer, 2021. 3(1): p. zcaa037.

23. Banchereau, R., et al., Personalized Immunomonitoring Uncovers Molecular Networks that Stratify Lupus Patients. Cell, 2016. 165(6): p. 1548–1550.

24. Davenport, E.E., et al., Discovering in vivo cytokine-eQTL interactions from a lupus clinical trial. Genome Biol, 2018. 19(1): p. 168.

25. Figgett, W.A., et al., Machine learning applied to whole-blood RNA-sequencing data uncovers distinct subsets of patients with systemic lupus erythematosus. Clin Transl Immunology, 2019. 8(12): p. e01093.

26. Guthridge, J.M., et al., Adults with systemic lupus exhibit distinct molecular phenotypes in a cross-sectional study. EClinicalMedicine, 2020. 20: p. 100291.

27. Hong, S., et al., Longitudinal profiling of human blood transcriptome in healthy and lupus pregnancy. J Exp Med, 2019. 216(5): p. 1154–1169.

28. Oon, S., et al., A potential association between IL-3 and type I and III interferons in systemic lupus erythematosus. Clin Transl Immunology, 2019. 8(12): p. e01097.

29. Toro-Domínguez, D., et al., Stratification of Systemic Lupus Erythematosus Patients Into Three Groups of Disease Activity Progression According to Longitudinal Gene Expression. Arthritis Rheumatol, 2018. 70(12): p. 2025–2035.

30. Dunning, J., et al., Progression of whole-blood transcriptional signatures from interferon-induced to neutrophil-associated patterns in severe influenza. Nature immunology, 2018. 19(6): p. 625–635.

31. Hoang, L.T., et al., Patient-based transcriptome-wide analysis identify interferon and ubiquination pathways as potential predictors of influenza A disease severity. PLoS One, 2014. 9(11): p. e111640.

32. Zhai, Y., et al., Host transcriptional response to influenza and other acute respiratory viral infections–a prospective cohort study. PLoS Pathog, 2015. 11(6): p. e1004869.

33. Le-Niculescu, H., et al., Towards precision medicine for stress disorders: diagnostic biomarkers and targeted drugs. Molecular Psychiatry, 2020. 25(5): p. 918–938.

34. Narang, V., et al., Influenza vaccine-induced antibody responses are not impaired by frailty in the community-dwelling elderly with natural influenza exposure. Frontiers in immunology, 2018. 9: p. 2465.

35. Rawat, C., et al., Downregulation of peripheral PTGS2/COX-2 in response to valproate treatment in patients with epilepsy. Scientific reports, 2020. 10(1): p. 1–14.

36. Tasaki, S., et al., Multi-omics monitoring of drug response in rheumatoid arthritis in pursuit of molecular remission. Nature communications, 2018. 9(1): p. 1–12.

37. Thakar, J., et al., Aging-dependent alterations in gene expression and a mitochondrial signature of responsiveness to human influenza vaccination. Aging (Albany NY), 2015. 7(1): p. 38.

38. Danon, L., et al., Comparing community structure identification. Journal of Statistical Mechanics: Theory and Experiment, 2005. 2005(09): p. P09008.

39. Rand, W.M., Objective criteria for the evaluation of clustering methods. Journal of the American Statistical association, 1971. 66(336): p. 846–850.

40. Levandowsky, M. and D. Winter, Distance between sets. Nature, 1971. 234(5323): p. 34–35.

41. Leek, J.T., et al., The sva package for removing batch effects and other unwanted variation in high-throughput experiments. Bioinformatics, 2012. 28(6): p. 882–3.

42. Risso, D., et al., Normalization of RNA-seq data using factor analysis of control genes or samples. Nature biotechnology, 2014. 32(9): p. 896–902.

43. Gerstner, J.R., et al., Removal of unwanted variation reveals novel patterns of gene expression linked to sleep homeostasis in murine cortex. BMC genomics, 2016. 17(8): p. 377–387.

44. Obeidat, M.e., et al., The effect of different case definitions of current smoking on the discovery of smoking-related blood gene expression signatures in chronic obstructive pulmonary disease. Nicotine & Tobacco Research, 2016. 18(9): p. 1903–1909.

45. NIH. Systemic Lupus Erythematosus (Lupus). 2020 [cited 2020 1st October]; Available from: https://www.niams.nih.gov/health-topics/lupus.

46. Buyon, J.P., et al., The effect of combined estrogen and progesterone hormone replacement therapy on disease activity in systemic lupus erythematosus: a randomized trial. Annals of internal medicine, 2005. 142(12_Part_1): p. 953–962.

47. Toro-Domínguez, D., et al., Stratification of systemic lupus erythematosus patients into three groups of disease activity progression according to longitudinal gene expression. Arthritis & Rheumatology, 2018. 70(12): p. 2025–2035.

48. Alhamdoosh, M., et al., Combining multiple tools outperforms individual methods in gene set enrichment analyses. Bioinformatics, 2017. 33(3): p. 414–424.

49. Munoz, L.E., et al., The role of defective clearance of apoptotic cells in systemic autoimmunity. Nature Reviews Rheumatology, 2010. 6(5): p. 280.

50. Banchereau, J. and V. Pascual, Type I interferon in systemic lupus erythematosus and other autoimmune diseases. Immunity, 2006. 25(3): p. 383–392.

51. Ivashkiv, L.B., IFNγ: signalling, epigenetics and roles in immunity, metabolism, disease and cancer immunotherapy. Nature Reviews Immunology, 2018. 18(9): p. 545–558.

52. Holland, S.M., Principal components analysis (PCA). Department of Geology, University of Georgia, Athens, GA, 2008: p. 30602–2501.

53. Borg, I. and P.J. Groenen, Modern multidimensional scaling: Theory and applications. 2005: Springer Science & Business Media.

54. Preparata, F.P. and M.I. Shamos, Computational geometry: an introduction. 2012: Springer Science & Business Media.

55. Kobak, D. and G.C. Linderman, UMAP does not preserve global structure any better than t-SNE when using the same initialization. bioRxiv, 2019.

56. Kriegel, H.-P., P. Kröger, and A. Zimek, Clustering high-dimensional data: A survey on subspace clustering, pattern-based clustering, and correlation clustering. ACM Transactions on Knowledge Discovery from Data (TKDD), 2009. 3(1): p. 1–58.

57. Butler, A., et al., Integrating single-cell transcriptomic data across different conditions, technologies, and species. Nature biotechnology, 2018. 36(5): p. 411–420.

58. Sun, S., et al., Accuracy, robustness and scalability of dimensionality reduction methods for single-cell RNA-seq analysis. Genome biology, 2019. 20(1): p. 269.

59. Dorrity, M.W., et al., Dimensionality reduction by UMAP to visualize physical and genetic interactions. Nature communications, 2020. 11(1): p. 1–6.

60. Sakaue, S., et al., Dimensionality reduction reveals fine-scale structure in the Japanese population with consequences for polygenic risk prediction. Nature communications, 2020. 11(1): p. 1–11.

61. Liberzon, A., et al., The molecular signatures database hallmark gene set collection. Cell systems, 2015. 1(6): p. 417–425.

