## Supplementary Table S2-S3, Figure S1-S7 for "Dimensionality reduction by UMAP reinforces sample heterogeneity analysis in bulk transcriptomic data"

Table S2: Dimensionality reduction methods, 4 methods

| Algorithm | Version | Availability | Implementation language | Parameter |
|---|---|---|---|---|
| UMAP | 0.3.10 | https://github.com/lmcinnes/umap/ | Python | n_components=2, n_neighbors=15, min_dist=0.1, metric='euclidean' |
| t-SNE | scikit-learn 0.23.1 | https://scikit-learn.org/ | Python | *n_components=2, perplexity=30.0, metric='euclidean', early_exaggeration=12.0, learning_rate=200.0, n_iter=1000,* |
| MDS | scikit-learn 0.23.1 | https://scikit-learn.org/ | Python | *n_components=2, *, metric=True, n_init=4, max_iter=300, verbose=0, eps=0.001, n_jobs=None, random_state=None, dissimilarity='euclidean'* |
| PCA | scikit-learn 0.23.1 | https://scikit-learn.org/ | Python | *n_components=2, copy=True, whiten=False, svd_solver='auto', tol=0.0, iterated_power='auto', random_state=None* |

Table S3: 5 clustering algorithms

| Algorithm | Version | Availability | Implementation language |
|---|---|---|---|
| k-means | scikit-learn 0.23.1: sklearn.cluster.KMeans | https://scikit-learn.org/ | Python |
| Hierarchical clustering | scikit-learn 0.23.1: sklearn.cluster.AgglomerativeClustering | https://scikit-learn.org/ | Python |
| Spectral clustering | scikit-learn 0.23.1: sklearn.cluster.SpectralClustering | https://scikit-learn.org/ | Python |
| gmm | scikit-learn 0.23.1: sklearn.mixture.GaussianMixture | https://scikit-learn.org/ | Python |
| hdbscan | hdbscan 0.8.18 | https://hdbscan.readthedocs.io/ | Python |

Figure S1: Clustering accuracy of five clustering algorithms on embedded space by four dimensionality reduction methods

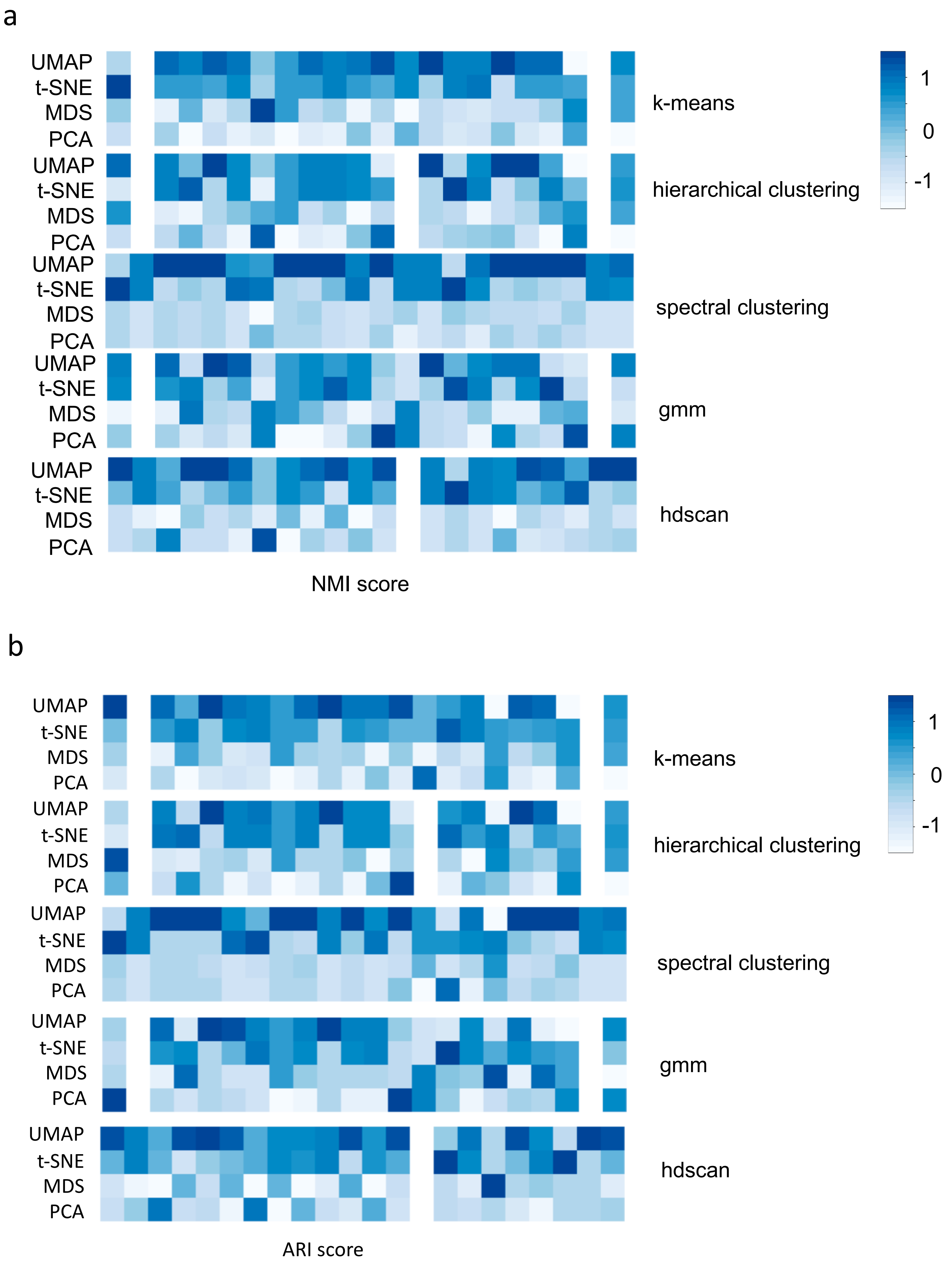

c

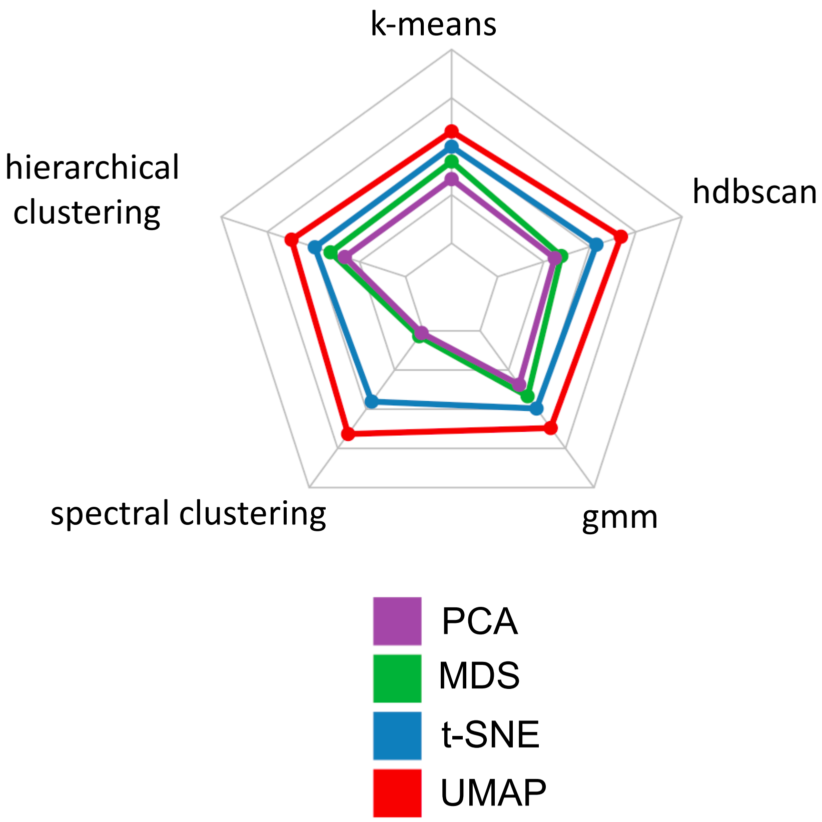


a. Heatmap for clustering accuracy by normalized mutual information (NMI). Darker blue means higher clustering accuracy. Five clustering algorithms are listed on the right side. [“1” in the heatmap represent the perfect accuracy, “0”…. “-1” represent….]
b. Heatmap for clustering accuracy by adjusted Rand index (ARI).
c. Radar plot of clustering accuracy (ARI score) comparison using five clustering methods. The average ARI score was on 22 datasets with cluster labels. The input was the embedded two-dimensional coordinates of each dimensionality reduction methods. Larger scale denotes better clustering accuracy, and UMAP outperformed the other three.

Figure S2: Neighborhood preserving (knn_k = 30), Average Jaccard index with 30 neighborhoods.

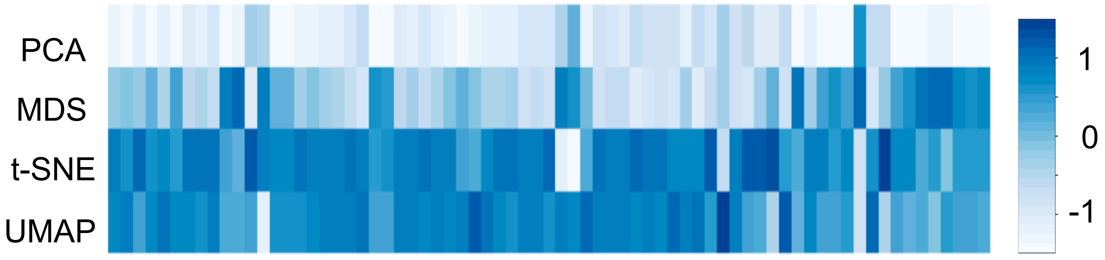


(***, N=71)

Heatmap for evaluating neighbourhood preserving of each method on 71 datasets. The number of neighbours is set as 30. The darker the colour is, the better the local information is retained.

Figure S3: Visualization of dataset GSE98793 and GSE107990 showing batch effects in two-dimensional space by dimensionality reduction methods.

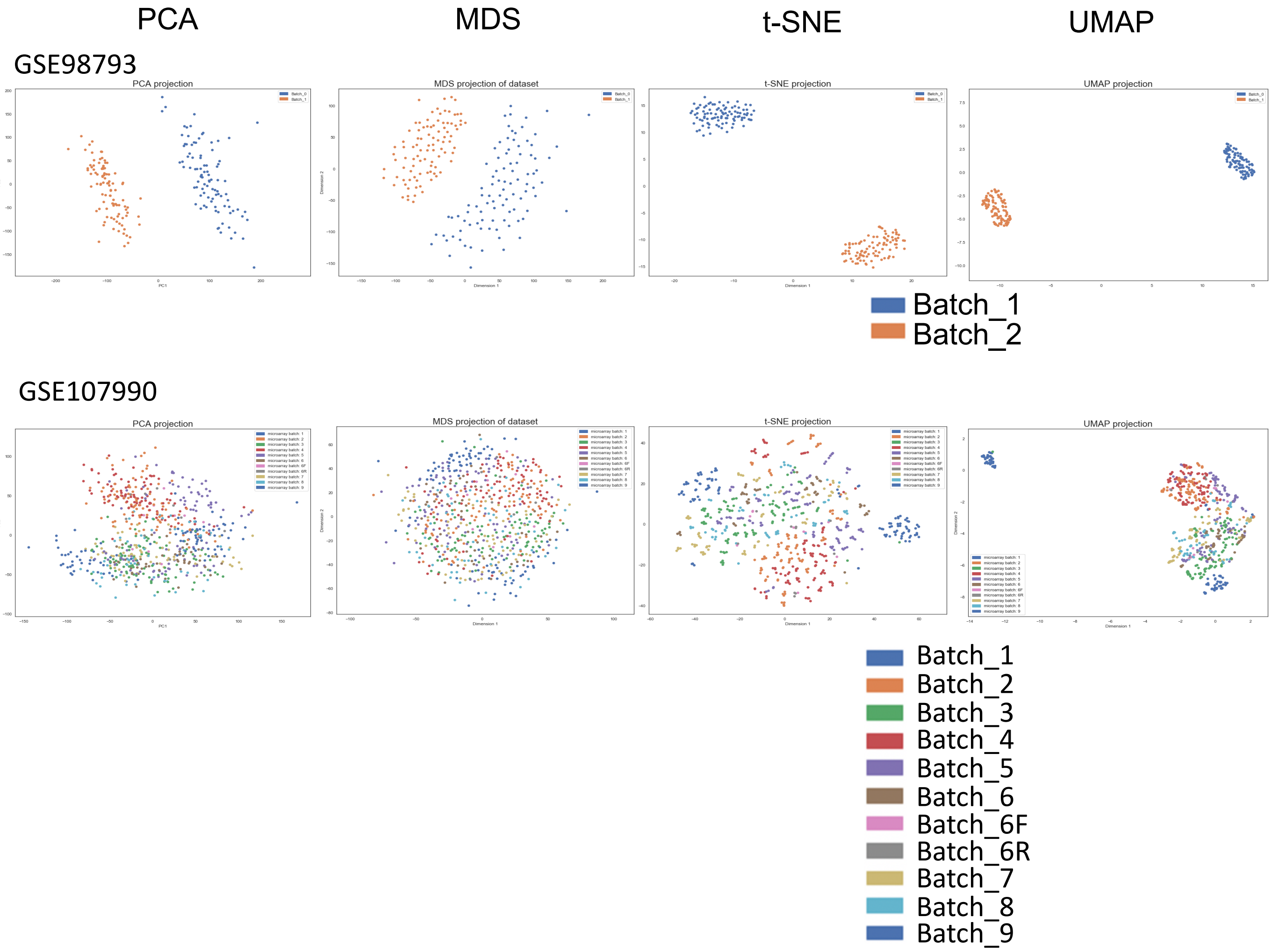

Figure S4: Visualization of eight datasets illustrating biological groups by dimensionality reduction methods. Colours represents groups.

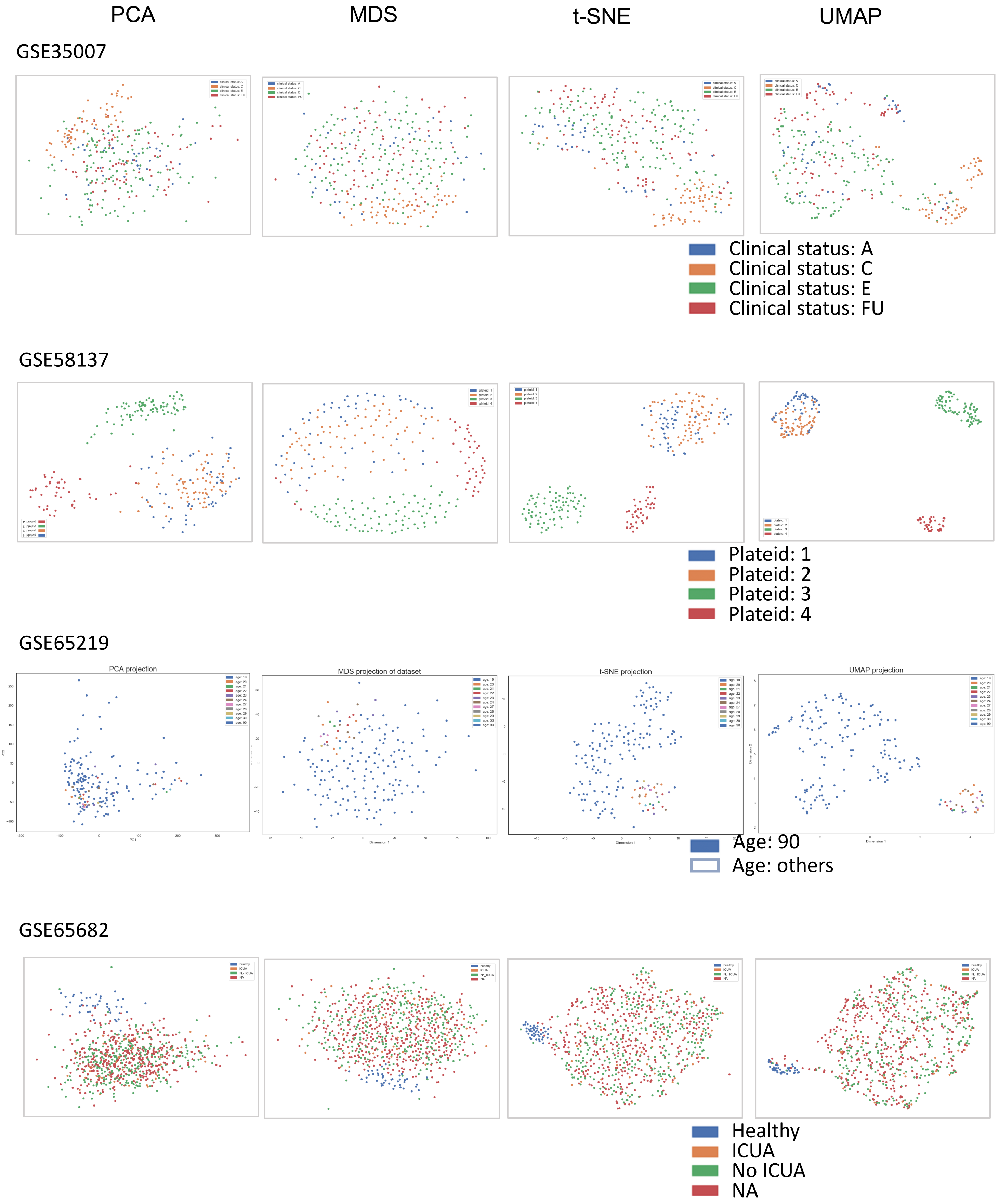

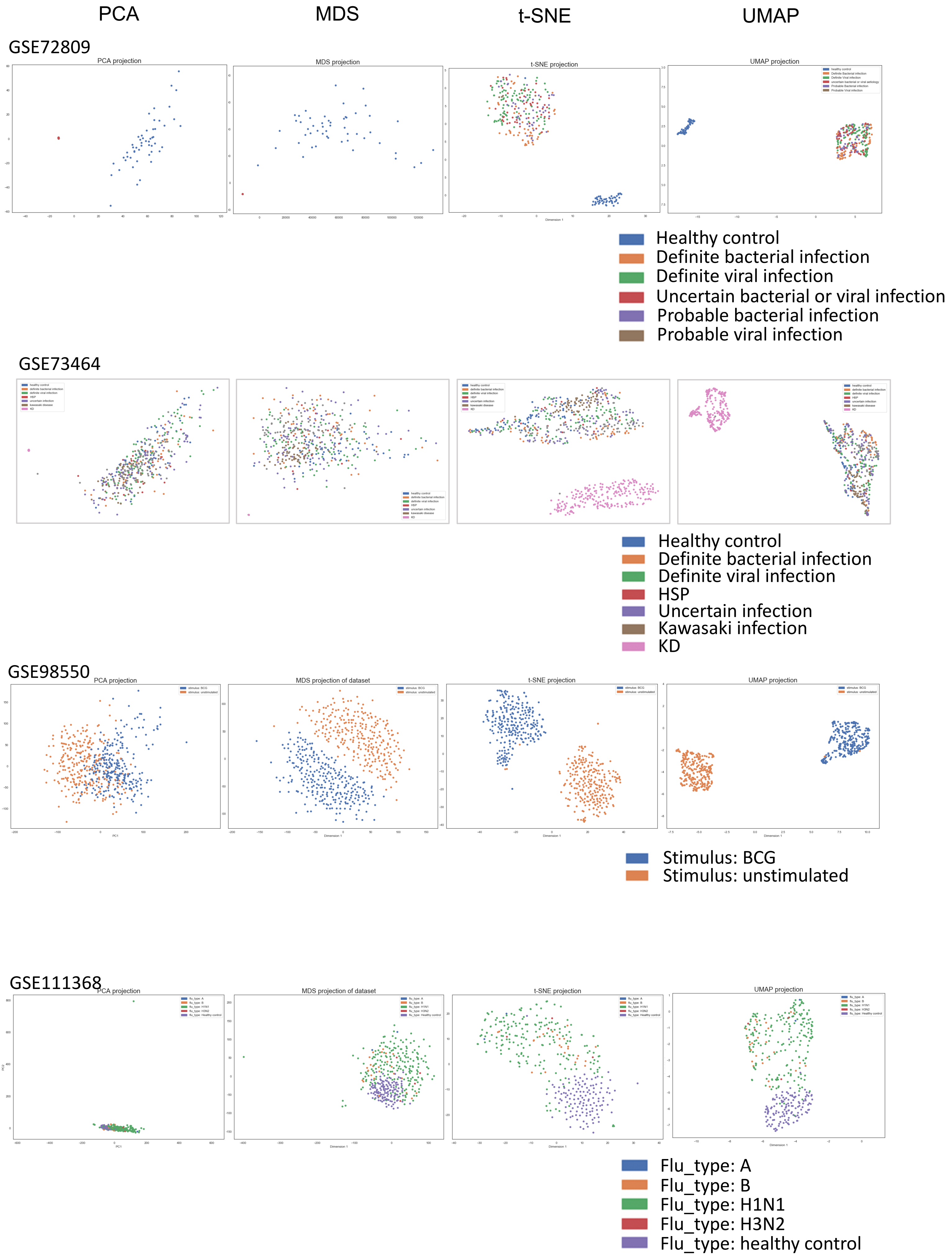

PCA
MDS
t-SNE
UMAP
GSE72809
PCA projection
MDS projection
t-SNE projection
UMAP projection
Healthy control
Definite bacterial infection
Definite viral infection
Uncertain bacterial or viral infection
Probable bacterial infection
Probable viral infection
GSE73464
Healthy control
Definite bacterial infection
Definite viral infection
HSP
Uncertain infection
Kawasaki infection
KD
GSE98550
PCA projection
MDS projection of dataset
t-SNE projection
UMAP projection
Stimulus: BCG
Stimulus: unstimulated
GSE111368
PCA projection
MDS projection of dataset
t-SNE projection
UMAP projection
Flu_type: A
Flu_type: B
Flu_type: H1N1
Flu_type: H3N2
Flu_type: healthy control

Figure S5: Associating sample features to clustering structure

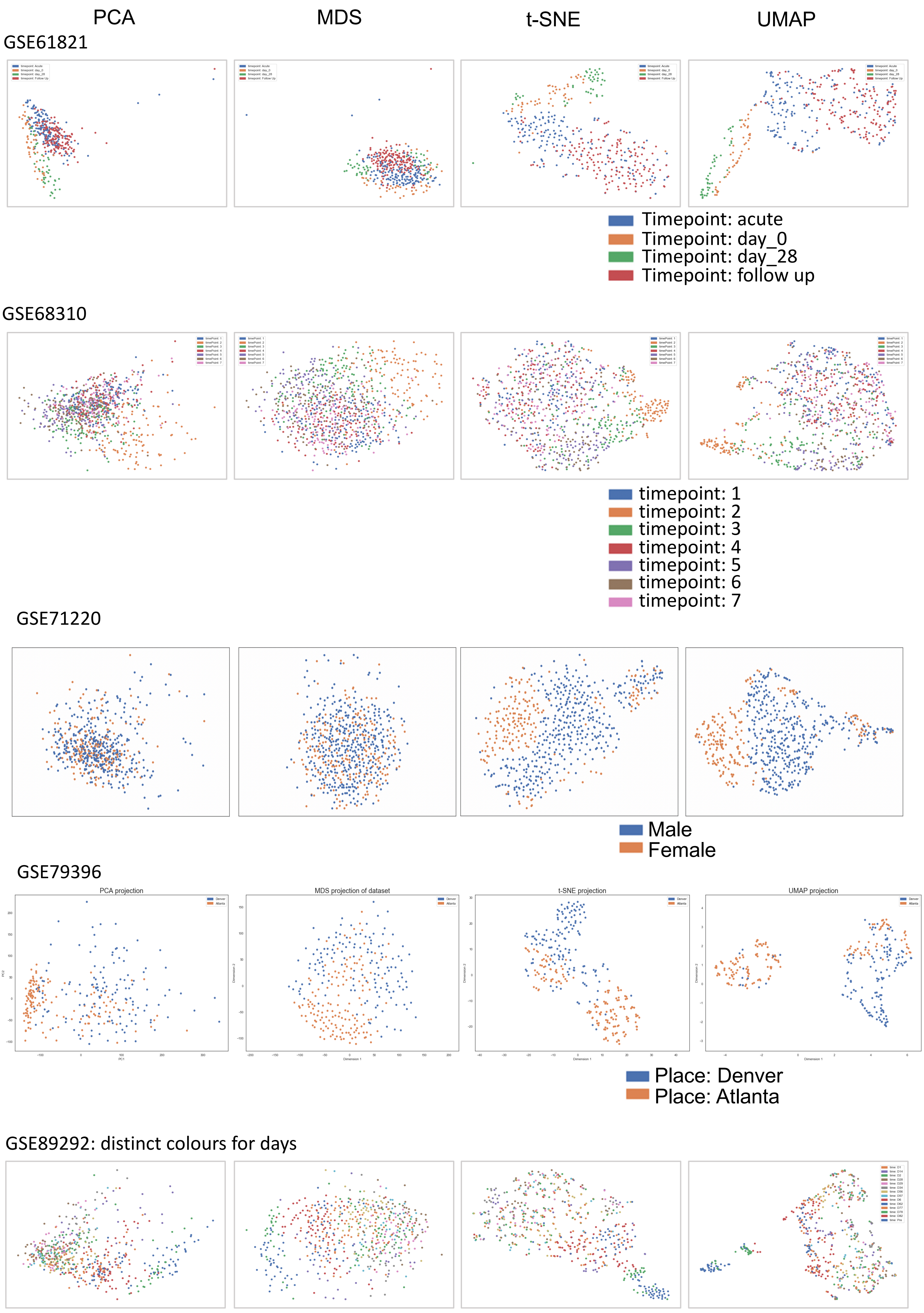

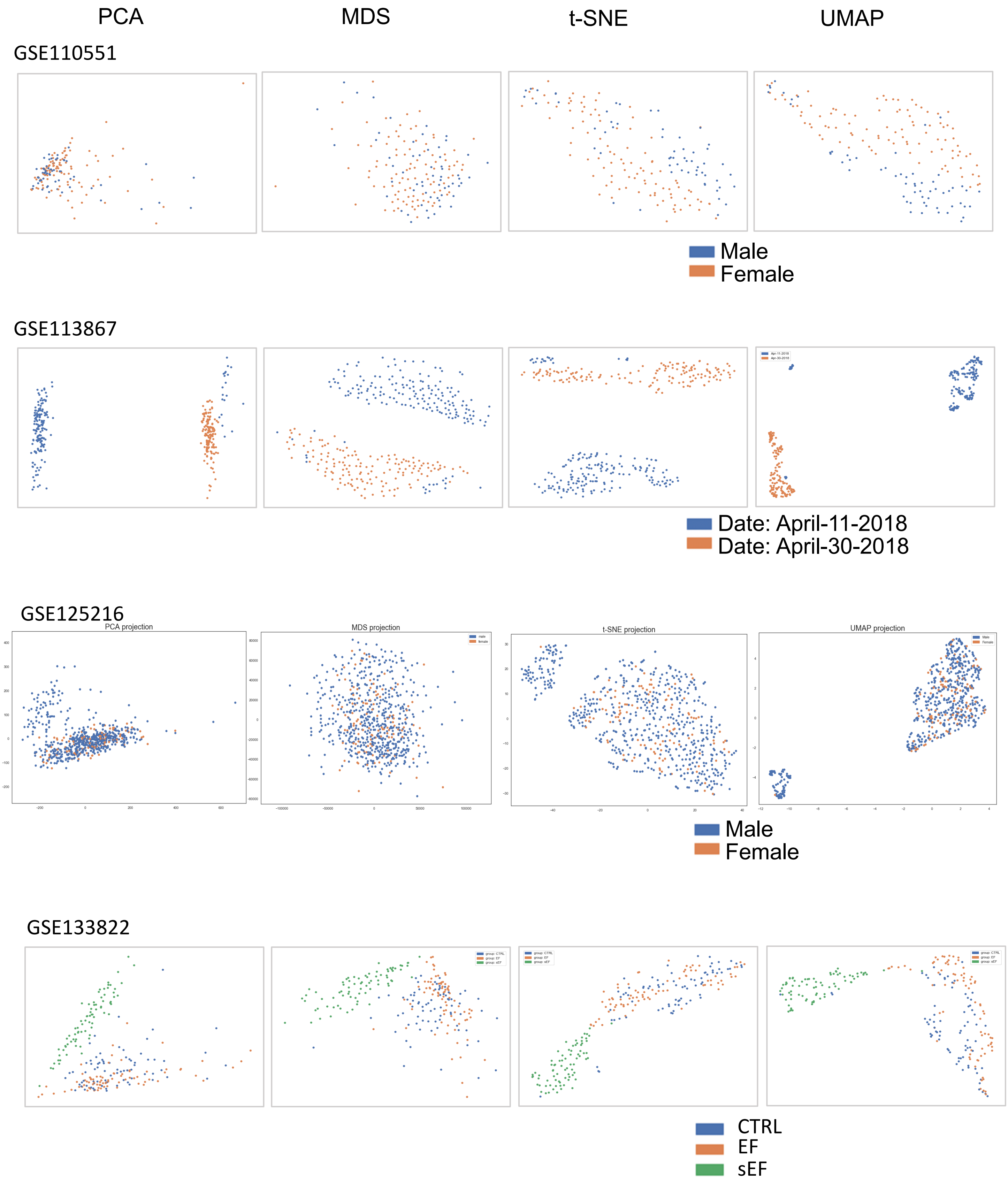
PCA
MDS
t-SNE
UMAP
GSE110551
Male
Female
GSE113867
Date: April-11-2018
Date: April-30-2018
GSE125216
PCA projection
MDS projection
t-SNE projection
UMAP projection
Male
Female
GSE133822
CTRL
EF
sEF

### Figure S6: Visualization of datasets showing clustering structures associated no pre-defined features

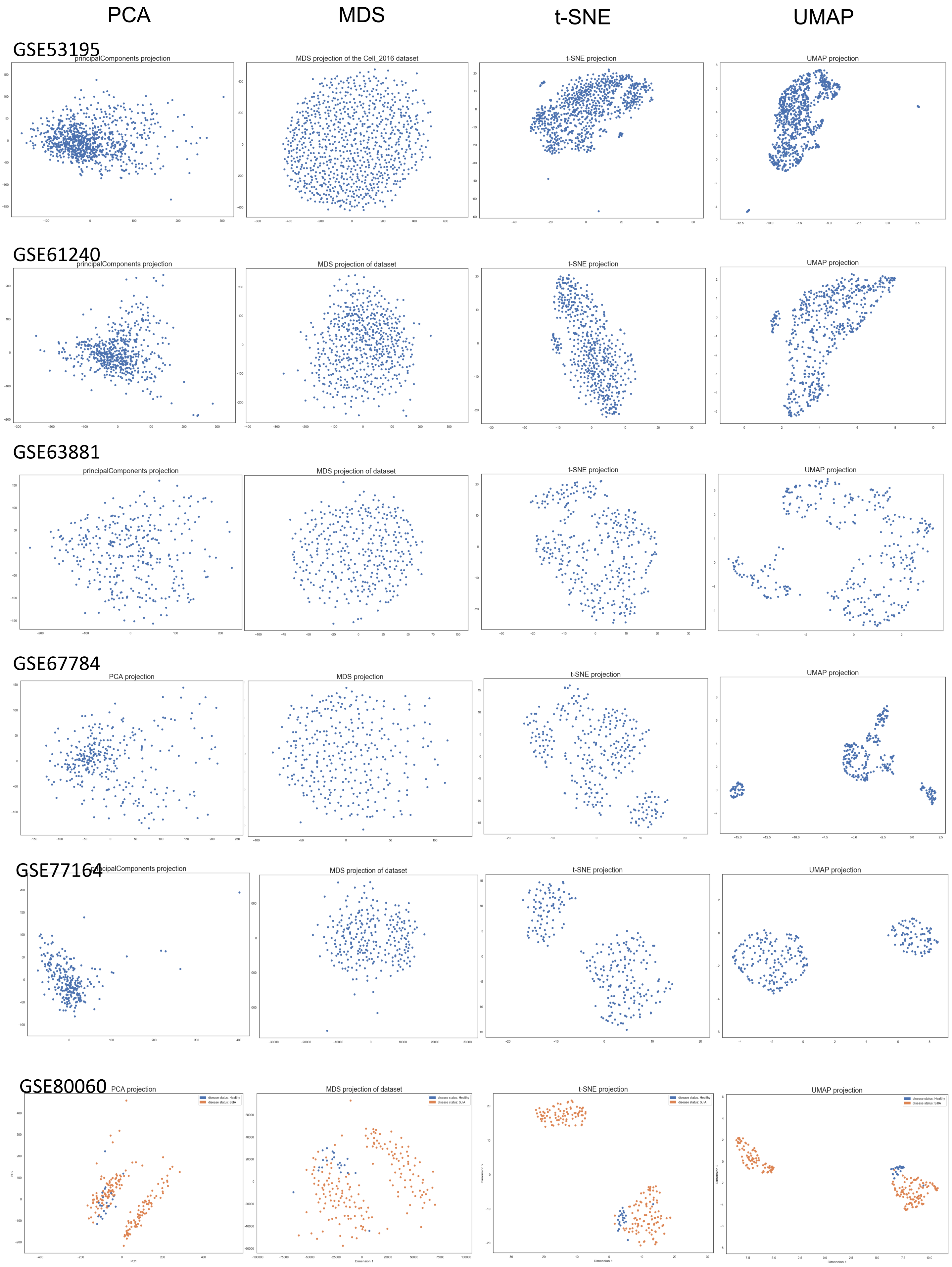

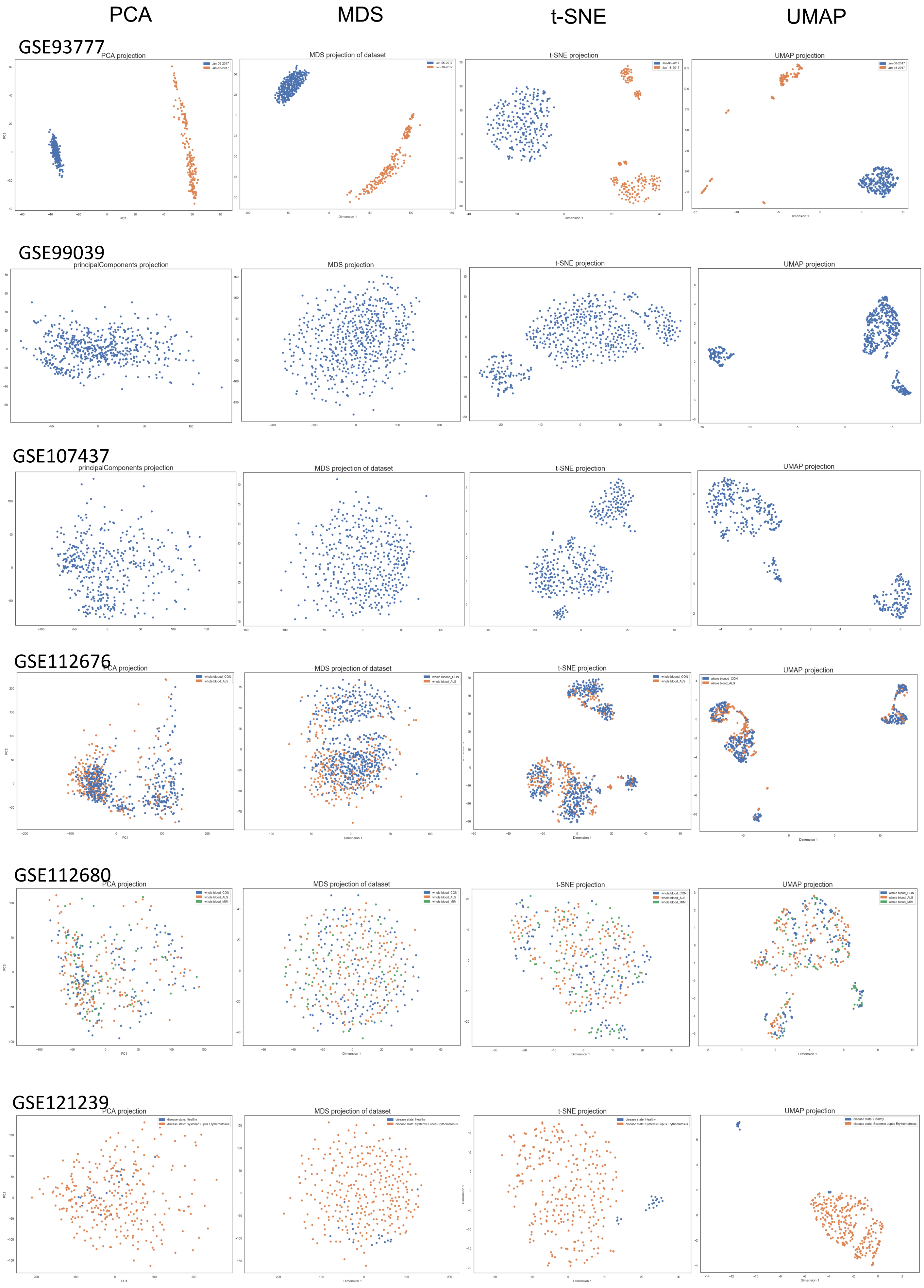
PCA
MDS
t-SNE
UMAP
GSE93777
PCA projection
MDS projection of dataset
t-SNE projection
UMAP projection
Jan-06-2017
Jan-18-2017
GSE99039
principalComponents projection
MDS projection
t-SNE projection
UMAP projection
GSE107437
principalComponents projection
MDS projection of dataset
t-SNE projection
UMAP projection
GSE112676
PCA projection
MDS projection of dataset
t-SNE projection
UMAP projection
whole bloood_CON
whole blood_ALS
GSE112680
PCA projection
MDS projection of dataset
t-SNE projection
UMAP projection
whole blood_CON
whole blood_ALS
whole blood_MIM
GSE121239
PCA projection
MDS projection of dataset
t-SNE projection
UMAP projection
disease state: Healthy
disease state: Systemic Lupus Erythematosus
PC1
PC2
Dimension 1
Dimension 2

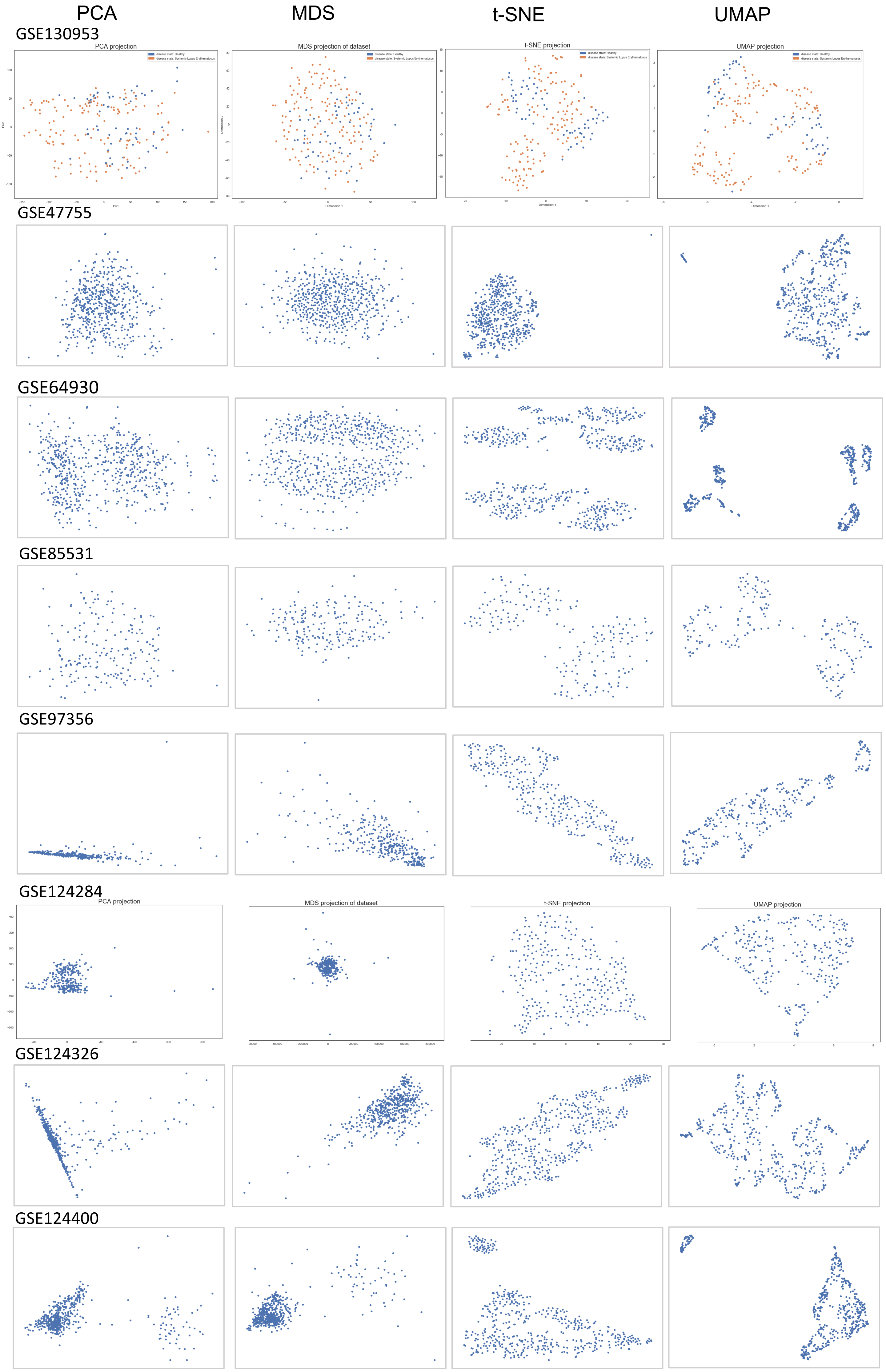
PCA
MDS
t-SNE
UMAP
GSE130953
PCA projection
MDS projection of dataset
t-SNE projection
UMAP projection
disease state: Healthy
disease state: Systemic Lupus Erythematosus
PC1
PC2
Dimension 1
Dimension 2
GSE47755
GSE64930
GSE85531
GSE97356
GSE124284
PCA projection
MDS projection of dataset
t-SNE projection
UMAP projection
GSE124326
GSE124400

Figure S7: Gene set enrichment analysis between sG1 v.s. sG0 and sG2 v.s. sG0 with top 20 differentially regulated molecular pathways ranked by adjusted p-value.

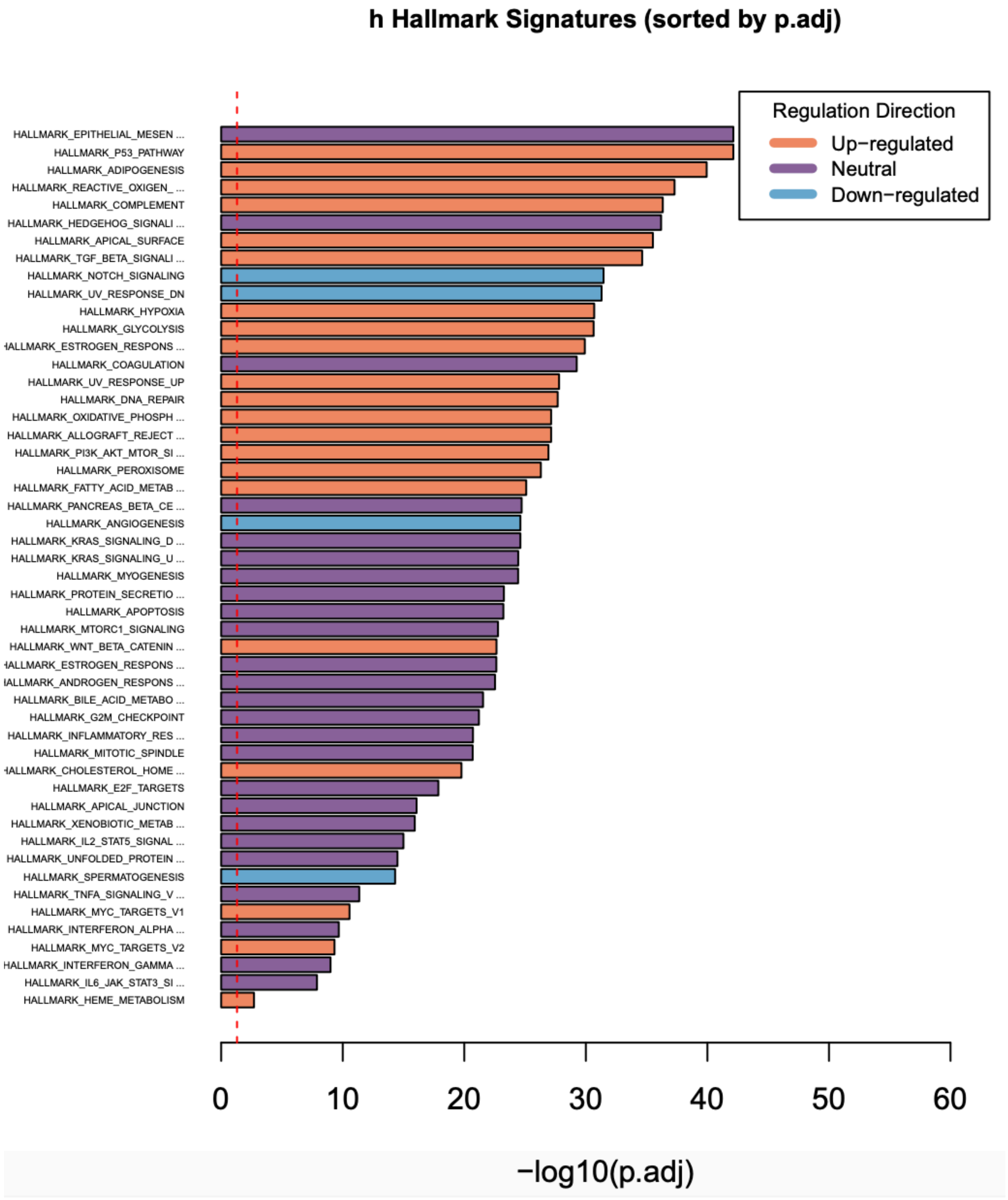
